# 3D Cell Neighbour Dynamics in Growing Pseudostratified Epithelia

**DOI:** 10.1101/2021.03.05.434123

**Authors:** Harold F. Gómez, Mathilde S. Dumont, Leonie Hodel, Roman Vetter, Dagmar Iber

**Author notes:** Corresponding author: Dagmar Iber.

## Abstract

During morphogenesis, epithelial sheets remodel into complex geometries. How cells dynamically organize their contact with neighbouring cells in these tightly packed tissues is poorly understood. We have used light-sheet microscopy of growing mouse embryonic lung explants, three-dimensional cell segmentation, and physical theory to unravel the principles behind 3D cell organization in growing pseudostratified epithelia. We find that cells have highly irregular 3D shapes and exhibit numerous neighbour intercalations along the apical-basal axis as well as over time. Despite the fluidic nature, the cell packing configurations follow fundamental relationships previously described for apical epithelial layers, i.e., Euler’s formula, Lewis’ law, and Aboav-Weaire’s law, at all times and across the entire tissue thickness. This arrangement minimizes the lateral cell-cell surface energy for a given cross-sectional area variability, generated primarily by the distribution and movement of nuclei. We conclude that the complex 3D cell organization in growing epithelia emerges from simple physical principles.

## INTRODUCTION

Common to all animals and plants, epithelia are a fundamental tissue type whose expansion, budding, branching, and folding is key to the morphogenesis of organs and body cavities. Characterized by apical-basal polarity (Figure 1a), epithelial cells adhere tightly to their apical neighbours in a virtually impermeable adhesion belt, form lateral cell-cell junction complexes along the apico-basal axis to provide mechanical stabilization, and bind tightly to the basal lamina and extracellular matrix (ECM) on the basal side (Drubin and Nelson, 1996; Rodriguez-Boulan and Macara, 2014; Shin and Margolis, 2006). How cell neighbour relationships are organised in these tightly adherent layers, and how these change during tissue and concomitant cell shape changes is poorly understood, despite their importance for cell-cell signalling and the fluidity of the tissue.

**Figure 1.**
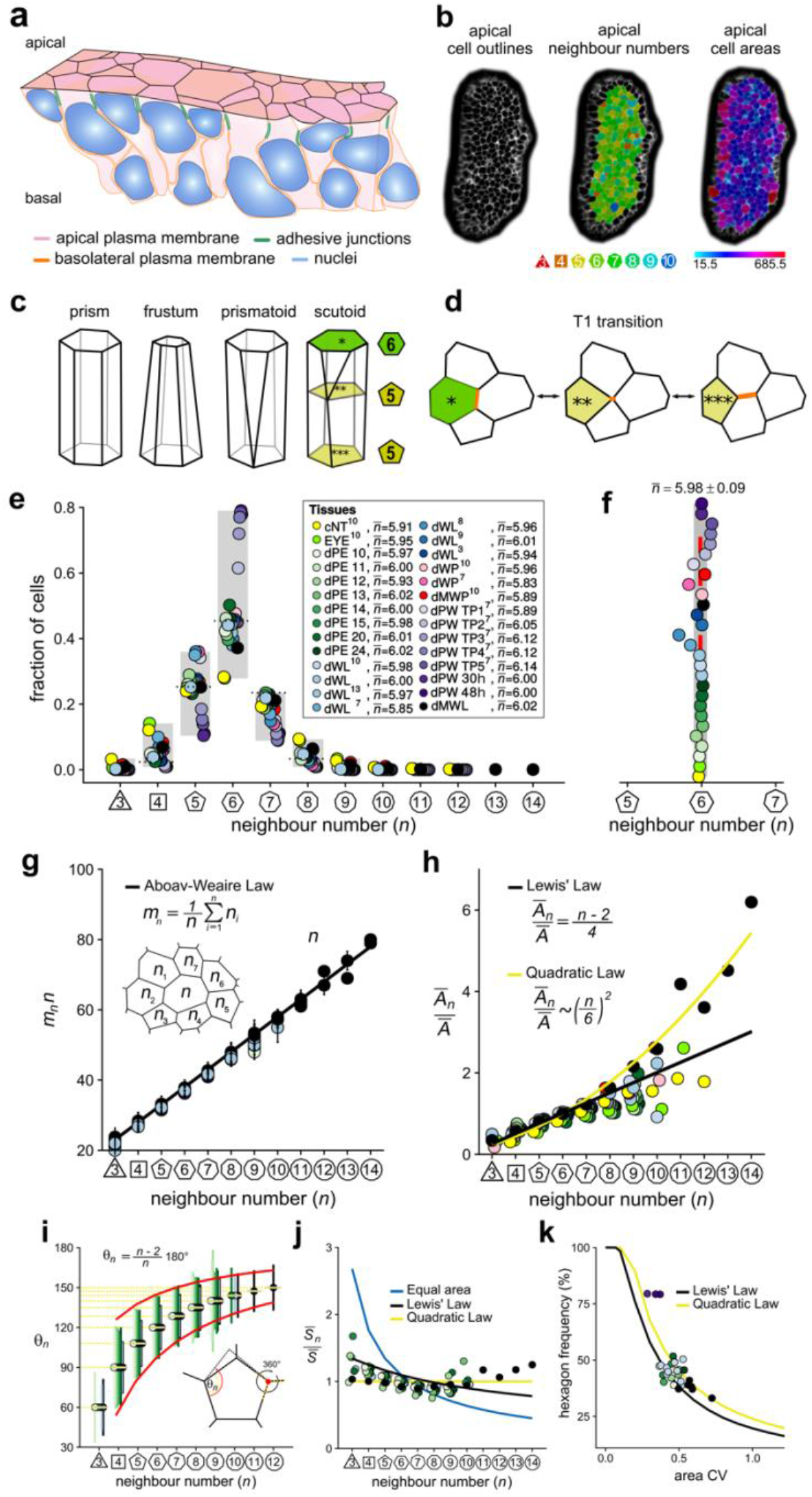
Principles of epithelial organization. **(a)** Schematic representation of an epithelial tissue layer. The cells are polarized between an apical and a basal side. Near the apical side, cells adhere tightly via adhesion junctions (green). Nuclei are depicted in blue. **(b)** Apical surface projection of an embryonic lung bud at E12.5 imaged using light-sheet microscopy. Cell contour segmentations (left) coloured according to neighbour relationships (middle) and area quantifications (right). **(c)** Current shape representations of 3D epithelial cells: prism, frustum, prismatoid, and scutoid. **(d)** Planar cell neighbour exchange (T1 transition). **(e)** Tissues differ widely in the frequency of neighbour numbers. The legend provides the measured average number of cell neighbours for each tissue and the references to the primary data (Classen et al., 2005; Escudero et al., 2011; Etournay et al., 2015; Farhadifar et al., 2007; Gibson et al., 2006; Heller et al., 2016; Sanchez-Gutierrez et al., 2016). Data points for n < 3 were removed as they must present segmentation artefacts. **(f)** The measured average number of cell neighbours is close to the topological requirement 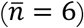 in all tissues; see panel e for the colour code. **(g)** Epithelial tissues follow the AW law (black line). The AW law formulates a relationship between the average number of neighbours, *n*, that a cell has and that its direct neighbours have, mn. The product mn.ncan be determined by summing over all *n*_*i*_. **(h)** The relative average apical cell area,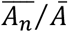, increases with the number of neighbours, *n*, and mostly follows the linear Lewis’ law (Eq. 2, black line), or the quadratic relationship (Eq. 3, yellow line) in case of higher apical area variability. **(i)** The average internal angle by polygon type is close to that of a regular polygon, *θn* = (*n*-2)/*n*.180° (yellow lines). To form a contiguous lattice, the angles at each tricellular junction must add to 360°, and the resulting observed deviation in the angles follows the prediction (red line). **(j)** The average normalised side length by polygon type. **(k)** Observed fraction of hexagons versus area coefficient of variation (CV). The curves mark theoretical predictions when polygonal cell layers follow either the linear Lewis’ law (Eq. 2, black line) or the quadratic law (Eq. 3, yellow line). The colour code in panels g-k is as in panel e, but data is available only for a subset of tissues. Panels e-k are reproduced with modifications from (Kokic et al., 2019) (Vetter et al., 2019).

Cell neighbour relationships can be most easily studied on epithelial surfaces, and the polygonal arrangements of apical surfaces (Figure 1b) have been meticulously analysed (Classen et al., 2005; Escudero et al., 2011; Etournay et al., 2015; Farhadifar et al., 2007; Gibson et al., 2006; Gómez-Gálvez et al., 2018; Heller et al., 2016; Kokic et al., 2019; Sanchez-Gutierrez et al., 2016). Widely considered to be a reliable proxy for three-dimensional (3D) cell shape, 3D epithelial cell shapes are often depicted as prisms with polygonal faces that retain the same neighbour relationships along the entire apico-basal axis (Figure 1c). Cells in curved epithelia are pictured as frustra, which have the same number of sides, but different apical and basal areas. If the curvature differs substantially along the principal axes, as is the case in epithelial tubes, neighbour relationships must change along the apical-basal axis. Prismatoids accommodate the neighbour change at the surface, while scutoids undergo the neighbour change somewhere along the apical-basal axis (Gómez-Gálvez et al., 2018). However, even though the curvature is the same in both principal directions of spherically shaped epithelia, the neighbour relationships still differ between the apical and basal sides (Gómez-Gálvez et al., 2018), suggesting that effects other than curvature must determine the 3D neighbour arrangements of cells in epithelia.

Given the challenges in visualising 3D neighbour arrangements, most studies to date have focused on apical cell arrangements, and have revealed striking regularities. First, even though the frequencies of neighbour numbers differ widely between epithelial tissues (Figure 1e), cells have on average (close to) six neighbours (Classen et al., 2005; Escudero et al., 2011; Etournay et al., 2015; Farhadifar et al., 2007; Gibson et al., 2006; Heller et al., 2016; Kokic et al., 2019; Sanchez-Gutierrez et al., 2016) (Figure 1f). This can be explained with topological constraints in contiguous polygonal lattices, as expressed by Euler’s Formula (Gibson et al., 2006; Rivier and Lissowski, 1982). Thus, if three cells meet at each vertex, the average number of neighbours in infinitely large contiguous polygonal lattices is exactly

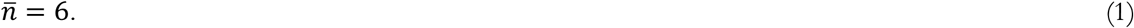

While the average number of neighbours in the entire lattice is (close to) to six, the local averages deviate from six, and instead rather closely follow a phenomenological relationship, termed Aboav-Weaire’s law (Aboav, 1970). According to Aboav-Weaire’s law (Figure 1g), the average number of neighbours of all *n* cells that border a cell with *n* neighbours follows as

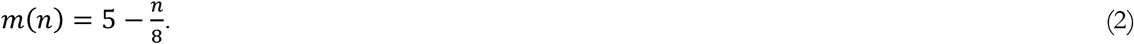

Finally, the average apical area, 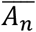, of cells with *n* neighbours is linearly related to the number of cell neighbours, *n* (Figure 1h, black line), a relation termed Lewis’ law (Lewis, 1928),

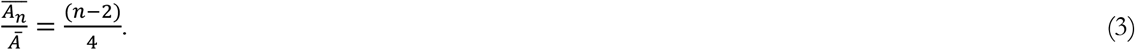

Here, ***Ā*** refers to the average apical cell area in the tissue.

We have recently shown that Aboav-Weaire’s law and Lewis’ law are a direct consequence of a minimisation of the lateral cell-cell contact surface energy (Kokic et al., 2019; Vetter et al., 2019). The lowest lateral cell-cell contact surface energy is obtained in a regular polygonal lattice because regular polygons have the smallest perimeter per polygonal area. The distribution of apical cell sizes that emerges from cell growth and division is, however, such that epithelial tissues cannot organise into perfectly regular polygonal lattices. By adhering to Aboav-Weaire’s law and Lewis’ law, cells assume the most regular lattice. In particular, by following, Aboav-Weaire’s law, the internal angles are closest to that of a regular polygon, while adding up to 360° at each tricellular junction (Figure 1i) (Vetter et al., 2019). And by following the relationship between polygon area and polygon type as stipulated by Lewis’ law (Eq. 2), the difference in side lengths, 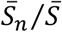, is minimized between cells (Figure 1j) (Kokic et al., 2019). The side lengths would be equal (Figure 1j, yellow line), if cells followed a quadratic relation of the form

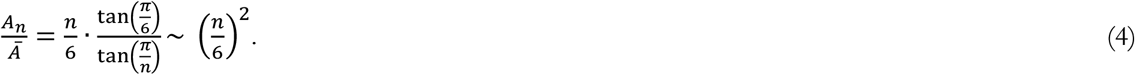

This quadratic relation (Figure 1h, yellow line), however, requires a larger area variability than is observed in most epithelia imaged to date. Accordingly, the predicted quadratic relation had not been previously reported, but could be confirmed experimentally by us by increasing the apical area variability (Kokic et al., 2019).

Given the relationship between apical area and neighbour numbers as stipulated by Eqs. 1,3,4, the apical area variability emerges as the key determinant of apical epithelial organisation, and the theory correctly predicts how the fraction of hexagons in the tissue depends on the apical area variability, as can be quantified by the coefficient of variation (CV = std/mean) (Figure 1k) (Kokic et al., 2019). As such, growth and cell division determine the variability of the apical areas and thus determine apical organisation indirectly. Taken together, the apical organisation of epithelia can be understood based on the principles of lateral cell-cell contact surface energy minimisation.

In this work, we leverage these theoretical insights along with light-sheet fluorescence microscopy to study 3D epithelial organisation, both in cleared and growing pseudostratified epithelia. We find that cells have complex 3D shapes with numerous neighbour transitions along their apical-basal axis as well as over time. We show that much as on the apical side, the variation of the cross-sectional areas along the apical-basal axis define the epithelial organisation at all times and across the entire tissue thickness. The observed neighbour arrangement minimizes the lateral cell-cell surface energy for a given cross-sectional area variability. The cross-sectional areas vary as a result of cell growth, division and interkinetic nuclear migration (IKNM). We conclude that the complex 3D cell organization in growing epithelia emerges from simple physical principles.

## RESULTS

### Apical and basal epithelial organisation

We started by exploring the apical and basal cellular organisation in epithelial tubes and buds (Figure 2a). To this end, we imaged CUBIC-cleared mouse embryonic (E12.5) lung rudiments from a Shh^GC/+^; ROSA^mT/mG^ background using light-sheet microscopy, and segmented the fluorescent membrane boundaries of over 400 cells per dataset in 2.5D (Figure 2b, Figure 2 -figure supplement 2). The apical and basal surfaces are both curved and thus differ in their total areas, i.e., the total segmented apical area is about 5-fold smaller than the basal area (Figure 2b). We detected less than half as many cells on the apical side, and the mean cross-sectional cell area of apical cells is therefore on average only 2-fold smaller than that of basal cells, while the area variability, measured as area CV, is higher (Figure 2c). Notably, the frequencies of the different neighbour numbers are not identical on the apical and basal side (Figure 2d), suggesting that the neighbour relationships change along the apical-basal axis, both in the tube and tip datasets. This observation is consistent with previous reports (Gómez-Gálvez et al., 2018). The change in neighbour relationships has previously been attributed to a curvature effect in tubes, but the neighbour changes in spherical geometries cannot be explained with such an effect (Gómez-Gálvez et al., 2018), suggesting that mainly other effects determine epithelial organisation.

**Figure 2.**
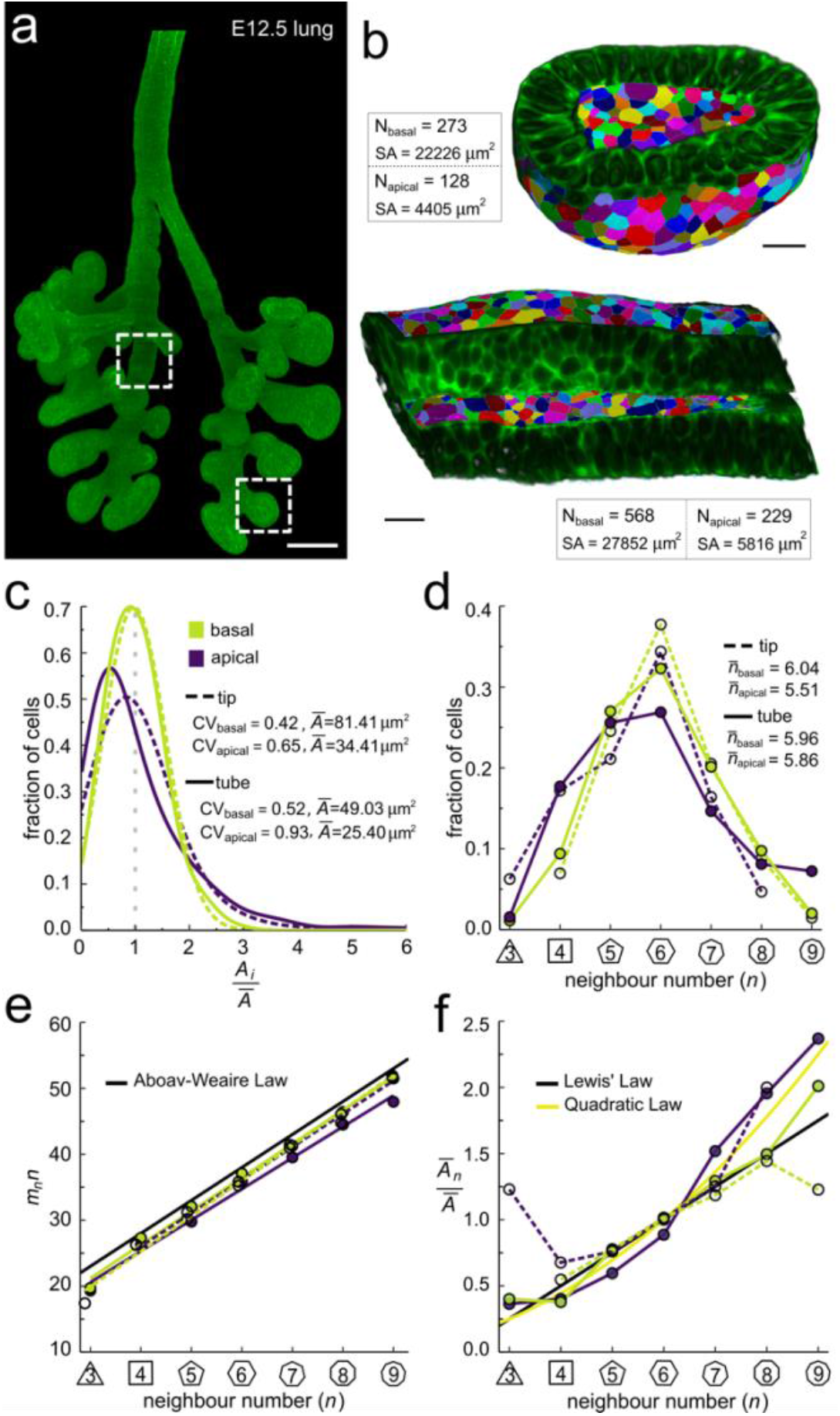
Apical and basal epithelial organization. **(a)** Epithelium of E12.5 ShhGC/+; ROSA^mT/mG^ mouse embryonic lung imaged using light-sheet microscopy. Scale bar 200 µm. Corresponding 2D sections are shown in Figure 2 -figure supplement 1, and Video 1.(b)Apical and basal 2.5D cell segmentation overlays on imaged tip and tube sections (dotted boxes in panel a). An illustration of the 2.5D segmentation workflow is presented in Figure 2 -figure supplement 2. Number of cells (N) and segmented surface areas (SA) are given. Cells are coloured using random labels. Scale bars 20 µm. **(c)** Normalised apical and basal cell area distributions in the tip (broken lines) and tube (solid lines) datasets. The colour code in panel c is reused in panels d-f. **(d)** Frequencies of neighbour numbers on the apical and basal sides in the tip and tube datasets. **(e)** The apical and basal layers follow the AW law (black line). **(f)** The normalised average cell area, 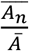, increases with the number of neighbours,*n*. The basal cells (green) follow Lewis’ law (Eq. 2, black line), while the apical cells (purple) follow the quadratic relationship (Eq. 3, yellow line).

So how can we explain the difference in apical and basal epithelial organisation in both datasets? We have previously shown that the apical organisation emerges from the minimisation of the overall lateral cell-cell contact surface energy (Kokic et al., 2019; Vetter et al., 2019). Aboav-Weaire’s law (Eq. 2, Figure 1g) and Lewis’ law (Eqs. 3,4, Figure 1h) emerge as global organization laws from this physical constraint, and ensure that the angles are closest to that of a regular polygon (Figure 1i, yellow lines), and that the side lengths are the most equal (Figure 1j, yellow line). We now find that Aboav-Weaire’s law (Figure 2e) and Lewis’ law (Figure 2f) hold not only for the apical, but also for the basal datasets. Consistent with our theory, the apical layers, which have a larger area variability than the basal layers (Figure 2c), follow the quadratic law (yellow line) rather than the linear Lewis’ law (black line).

We conclude that basal layers follow the same organisational principles as apical layers, such that their organisation can also be explained with a minimisation of the lateral cell-cell contact surface energy. Accordingly, the observed difference in overall neighbour relationships (Figure 2e,f) is a consequence of the difference in the cross-sectional area distributions (Figure 2c). So, why do the normalised area distributions differ between the apical and basal sides in both the tube segment and the bud, and how do they change along the apical-basal axis?

### 3D organisation of epithelia

To explore the physical principles behind 3D epithelial cell organisation, we 3D segmented 140 cells from a tube segment and 59 cells from a bud segment in CUBIC-cleared, light-sheet imaged embryonic lung explants (Figure 3a, Figure 2 -figure supplement 1, Video 1-4). By interpolating between equally spaced sequential contour surfaces (every 1.66 µm in the tube and every 1.72 µm in the bud dataset) along the apical-basal axis, accurate volumetric reconstructions of cell morphology were obtained that allowed for the extraction of morphometric quantifications along the apical-basal axis. In both datasets, the 3D organisation of epithelial cells is highly complex, and cell neighbour relationships change continuously along the apical-basal axis (Figure 3b). As a result, cells are in direct physical contact not only with the cells that are neighbours on the apical side, but also with cells that appear two or even three cell diameters apart (Figure 3c).

**Figure 3.**
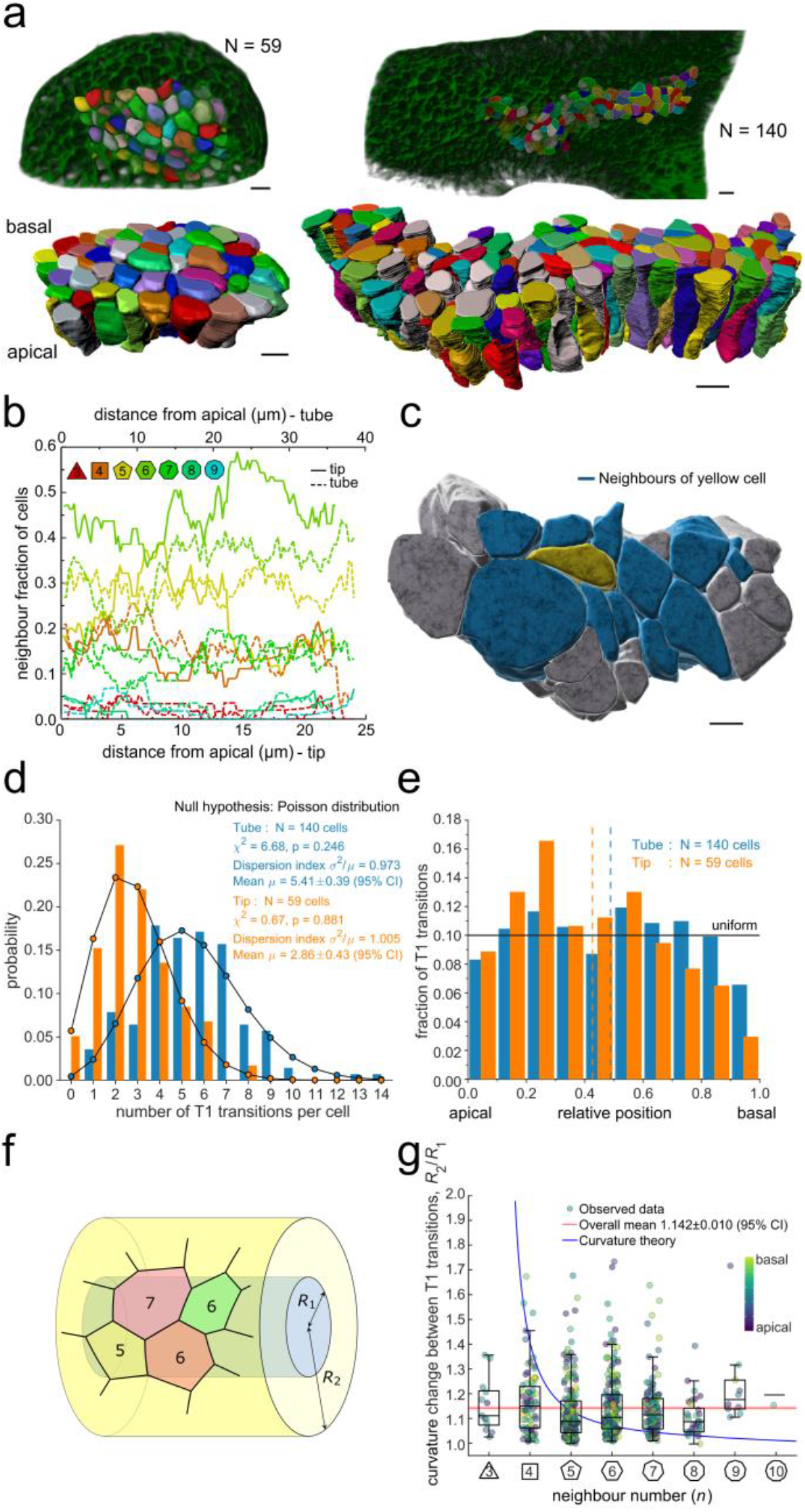
3D Epithelial Organisation. **(a)** 3D iso-surfaces of segmented epithelial tip (N=59) and tube (N=140) cells from a Shh^GC/+^; ROSA^mT/mG^ E12.5 mouse lung rudiment imaged using light-sheet microscopy. Morphometric quantifications of cell boundary segmentations along the apical-basal axis were used to study spatial T1 transitions. The 3D segmentation workflow is introduced in Figure 3 -figure supplement 1 and Video 2-4 illustrate the rendered epithelial tip, tube and all segmented volumes. Scale bars 10 µm. **(b)** Frequency of neighbour numbers as quantified along the apical basal axis in the tip (solid lines) and tube (broken lines) datasets. **(c)** Extent of neighbour contacts (center cell in yellow and neighbours in blue) in 3D as viewed from the apical side. Scale bars 5 µm. **(d)** Probability distributions of the lateral T1 transitions for tip (total=169, mean=2.86, N=59) and trunk (total=746, mean=5.41, N=140) datasets are consistent with Poisson distributions. **(e)** Normalized apical-basal distribution of T1 transitions for all cells shows no apical-basal bias, except for fewer transitions close to the basal surface. **(f)**. Schematic of a tubular epithelium. Along the apical-basal axis, the tissue curvature reduces from 1/*R*_1_ to 1/*R*_2_. **(g)** The predicted impact of a curvature effect on T1 transitions decreases with increasing cell neighbour numbers (blue line). The measured T1 transitions for different neighbour numbers do not support a curvature effect (dots, boxplots, and red line). Boxplots indicate the median, 25% and 75% percentiles of the data.

Remarkably, we record up to 14 cell neighbour changes per cell in the tube and up to 8 in the tip, between adjacent cross-sections along the apical-basal cell axis (Figure 3d). We will refer to these neighbour changes as lateral T1 transitions, or T1L. The mean relative apical-basal position for the lateral T1 transitions is 0.489 ±0.020 (95% CI), and there is no clear apical or basal tendency, though fewer transitions are observed close to the basal surface (Figure 3e). The dispersion index, i.e. the ratio of the variance σ^2^ and the mean number µ of transitions per cell, which equals unity for a Poisson distribution, is close to unity for both samples (Figure 3d). The chi-squared test also confirms that the number of apical-basal T1 transitions per cell is Poisson-distributed (Figure 3d). A Poisson distribution models the probability of a number of independent random events occurring in a given interval at a constant average rate. The consistency with a Poisson distribution, therefore, suggests a stochastic basis to the 3D organization of epithelial cells.

The large number of observed T1L transitions and their distribution along the apical-basal axis challenges the recently popularized notion of curvature-driven scutoids as cell building blocks for epithelia (Gómez-Gálvez et al., 2018). To further examine the potential influence of tissue curvature on T1L transitions, we measured the apical-basal distance between two consecutive neighbour number changes for each cell in the tube dataset and recorded at which local tissue curvature they occur. For this analysis, we excluded cell portions from the apical end to the first transition and from the last transition to the basal end to reduce boundary effects, i.e., only interior segments between transitions were considered. The mean apical-basal distance between two transitions is 17.89 ± 0.66 µm (95% CI). Local tissue curvature was approximated by fitting ellipses to the apical and basal surface boundaries of the tubular epithelium in 624 equidistant sections perpendicular to the main tube axis. The semi-axes of these ellipses were then averaged over all sections to obtain the semi-axes *a*_apical_, *a*_basal_, *b*_apical_, *b*_basal_. Since our sample of 140 cells was segmented from a region close to the cusp of the nearly elliptical tube, a reasonably close estimate of the local tissue curvature where a T1L transition occurs is given by a linear interpolation between the curvature at the minor vertices of the apical and basal ellipses, according to the relative apical-basal position of the transition. The minor curvature of an ellipse with major and minor semi-axes *a* and *b* is given by *b/a*^*2*^. Therefore, we estimate the local radius of curvature *R* by

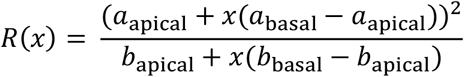

where *x*ϵ[0,1] is the relative apical-basal location of the T1L transition. The examined tissue exhibits an average curvature fold change of *R(1)/R(0)*=2.21 from the basal to the apical side. Denoting by R_1_ and R_2_ the radii of curvature between two adjacent T1 transitions along the apical-basal axis of a cell (Figure 3f), we find that the distribution of curvature fold change *R*_2_/*R*_1_ shows no significant dependency on the number of neighbours *n* the cell has along that portion of the cell (Figure 3g). The mean curvature fold change per apical-basal T1L transition per cell is <R_2_/R_1_> = 1.142 ± 0.010 (95% CI). By extending the theory of scutoids (Gómez-Gálvez et al., 2018) to multiple T1L transitions per cell, we have derived a quantitative estimate of how tissue curvature would translate into the number of neighbour exchanges within that framework (Supplementary Material). If curvature changes were a main driver of T1L transitions, cells with smaller neighbour numbers n would be expected to change n over a much larger curvature fold change than cells with many neighbours (Figure 3g, blue line). However, we observe no systematic dependency of the curvature fold change on the number of neighbours the cell has along that portion of the cell in the developing mouse lung epithelium (Figure 3g). From this, we conclude that tissue curvature affects cell neighbourhood rearrangements through the tissue thickness at most mildly.

### Neighbour changes along the apical-basal axis are driven by changes in cross-sectional area variation

Other than curvature effects, what else could drive the observed changes in neighbour relationships along the apical-basal axis? We notice that much as the apical and basal layers, each layer along the apical-basal axis behaves according to the three relationships previously described for the apical side, i.e., Euler’s formula (Eq. 1, Figure 4a), Aboav-Weaire’s law (Eq. 2, Figure 4b), and Lewis’ law (Eqs. 3, 4, Figure 4c). As predicted by the theory based on the minimisation of the lateral cell-cell energy (Kokic et al., 2019), the layers with a large area variability (Figure 4a) follow the quadratic law (yellow line) and those with a lower area variability the linear Lewis’ law (black line). The fraction of hexagons also follows the predicted relationship with the cross-sectional area variability (Figure 4d). We conclude that the entire 3D organisation of epithelia can be explained with a minimisation of the lateral cell-cell contact surface energy, as previously revealed for the apical layer.

**Figure 4.**
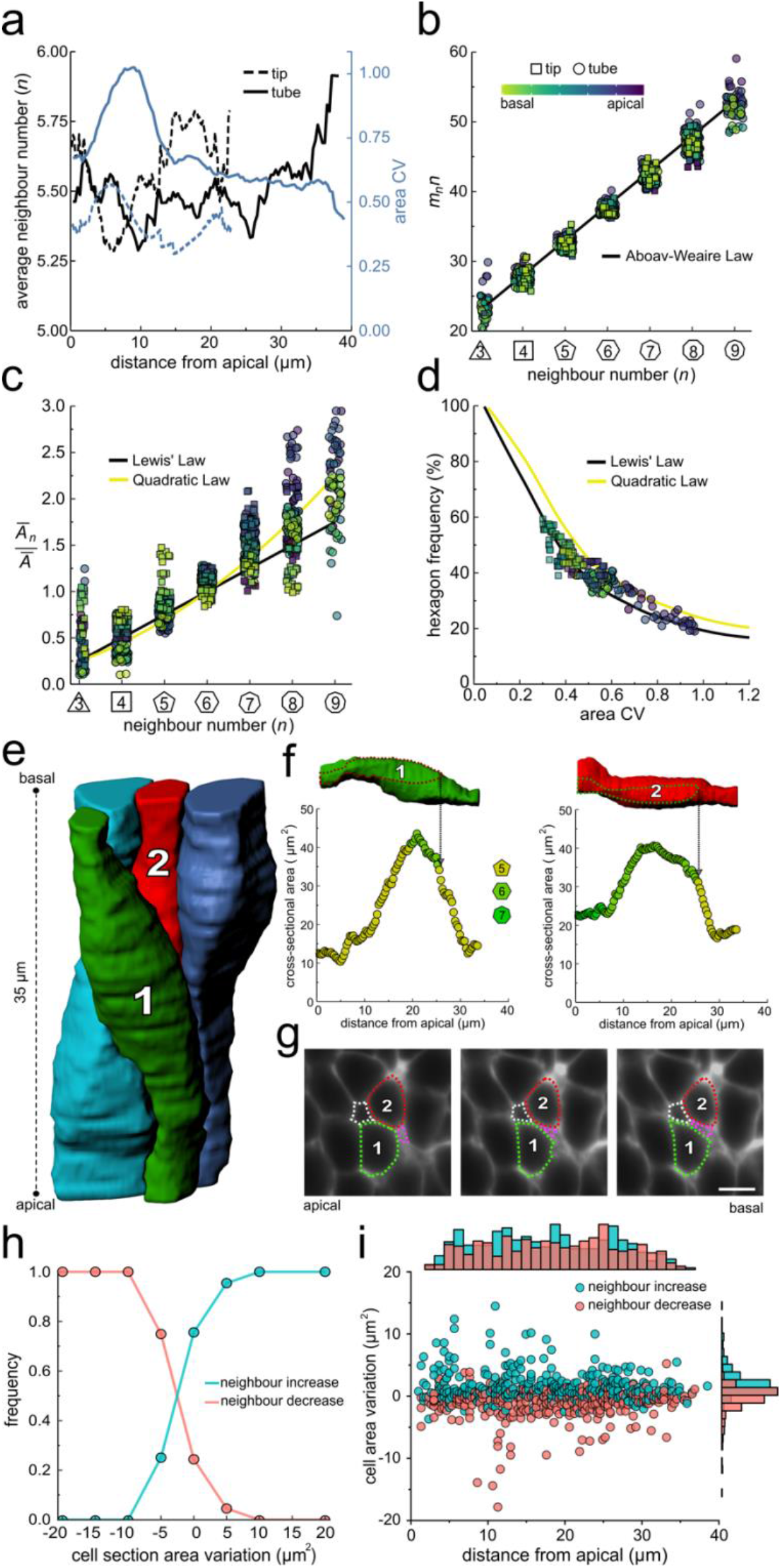
Neighbour changes along the apical-basal axis are driven by changes in cross-sectional area. **(a)** Average number of neighbours (black) and area CV (blue) along the apical basal axis in the tip and tube datasets. **(b)** All epithelial layers follow the AW law (black line). The colour code in panels c, d follow that in panel b. **(c)** All epithelial layers follow Lewis’ law (Eq. 2, black line) in case of low, and the quadratic relationship (Eq. 3, yellow line) in case of high cell area variability. **(d)** Observed fraction of hexagons versus area CV for segmented cell layers along the apical-basal axis. The lines mark the theoretical prediction if polygonal cell layers follow either the linear Lewis’ law (black line) or the quadratic law (yellow line). **(e)** 3D iso-surfaces of four segmented epithelial cells in a CUBIC-cleared Shh^GC/+^; ROSA^mT/mG^ E12.5 distal lung tube, with 140 3D segmented epithelial cells (Figure 3 -figure supplement 1c). **(f)** Cross-sectional area and cell neighbour number along the apical-basal axis for marked cells in panel e. Dotted lines indicate contact with one another. **(g)** Lateral cross-sections illustrating a T1 transition along the apical-basal axis (0.664 µm in-between frames). Scale bar 6 µm. **(h)** An increase in the cell cross-sectional area increases the frequency of a neighbour number increasing spatial T1 transitions, and vice versa. **(i)** Apical-basal distribution of spatial T1 transitions according to neighbour increase or decrease and cross-sectional area variation.

If epithelial cell neighbour relationships are indeed driven by a minimisation of the total lateral cell-cell contact surface energy, then the T1L transitions along the apical-basal axes should be driven by changes in the cross-sectional area along the apical-basal axis. If we analyse four 3D segmented cells (Figure 4e) in detail, we indeed see how an increase in the cross-sectional area results in an increase in the neighbour number, and *vice versa* (Figure 4f) via lateral T1 transitions (Figure 4g). As the cell neighbour arrangements represent global minima, the local analysis does, of course, not provide a perfect correlation. When we consider all 140 segmented cells in the tube segment with their 746 cell neighbour exchanges between adjacent cross-sections (Figure 3e), then we find that the frequency of T1L transitions along the apical-basal axis is indeed higher, the larger the increase in cross-sectional area, and vice versa (Figure 4h,i).

### Changes in cross-sectional area as a result of interkinetic nuclear migration (IKNM)

So, what determines the cross-sectional cell areas in each layer? In epithelia, mitosis is restricted to the apical surface. Depending on the average diameter of nuclei and the average apical cross-sectional area, there is insufficient space for all nuclei to be accommodated apically. Therefore, as a cell exits mitosis, the nucleus moves from the apical towards the basal side (G1 and S phase) and then back to the apical side (G2 phase) to undergo another round of mitosis, a process referred to as interkinetic nuclear migration (IKNM) (Meyer et al., 2011). Consequently, nuclei are distributed along the entire apical-basal axis, giving the tissue a pseudostratified configuration. We wondered to what extent the nuclear distribution, and its effect on the 3D cell shape, explains the observed area distributions and lateral T1 transitions.

To this end, we stained the nuclear envelope with fluorescently tagged antibodies against lamin B1 (Figure 5a, Figure 5 -figure supplement 1), and 3D segmented all nuclei within epithelial cells in a tube segment (Figure 5b, Figure 3 -figure supplement 1, Video 3). The nuclei were distributed along the entire apical-basal axis (Figure 5c), and consistent with the pseudostratified appearance of the epithelium, nuclei in neighbouring cells had different positions along the apical-basal axis (Figure 5b). The nuclear shapes, volumes, and cross-sectional areas (Figure 5d-f) all varied along the apical-basal axis. As expected, nuclei are largest and most spherical at the apical side, where they undergo mitosis (Figure 5d). Thus, a one-sided, two-sample Welch t-test revealed a significantly reduced ellipticity of nuclei located in the first 25% of the apical-basal axis compared to those in the middle 50% (p=0.0002). While the nuclear volumes (Figure 5f) and cross-sectional areas (Figure 5g) are slightly smaller than for the entire cell, the cross-sectional areas of the cell and the nucleus are strongly correlated (r = 0.94, Figure 5g). This is consistent with the nucleus determining the cross-sectional cell area, where present. Cell sections without nucleus typically have smaller cross-sectional areas, thereby leading to a higher frequency of small cross-sections in cells compared to nuclei. Accordingly, as previously seen for the cell cross-sectional areas, the observed changes in cell neighbour numbers correlates with the observed changes in nuclear cross-sectional areas (Figure 5h) such that most T1L transitions occur where the nucleus starts and ends (Figure 5i).

**Figure 5.**
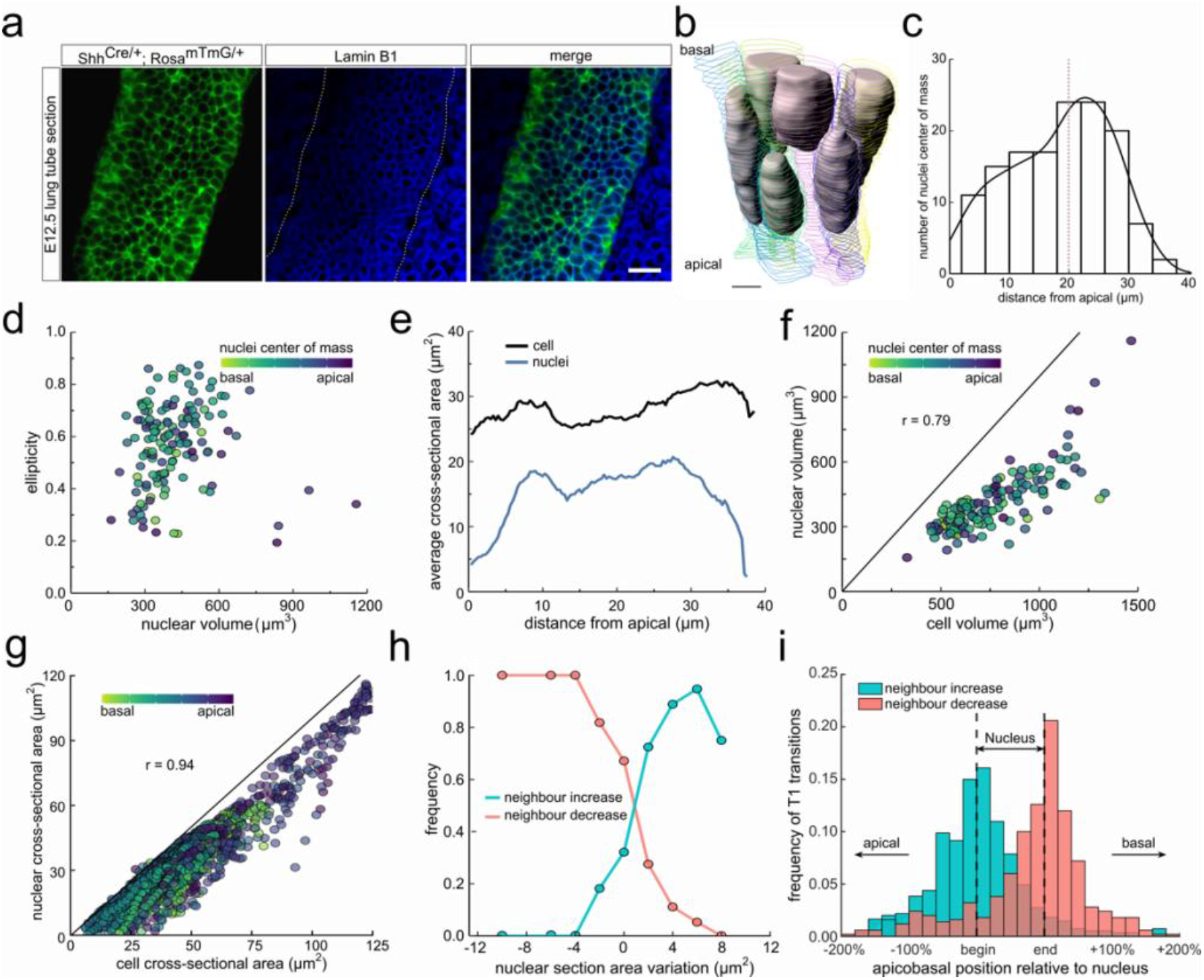
Changes in cross-sectional area as a result of interkinetic nuclear migration (IKNM). **(a)** Light-sheet microscopy longitudinal sections of an E12.5 CUBIC-cleared lung tube carrying the Shh^GC/+^; ROSA^mT/mG^ reporter allele (green epithelium) and immunostained for lamin B1 (blue nuclear envelopes). Morphometric quantifications of 3D iso-surfaces (N=140) and cell segmentations along the apical-basal axis were used to study the nature of cross-sectional area variation and the effect of IKNM (Figure 3 -figure supplement 1). Scale bar 20 µm. **(b)** Sequential cell membrane contour surfaces and nuclear iso-surfaces for six epithelial cells. By interpolating between contours and creating iso-surfaces, 3D shapes can be accurately extracted (Video 3,5). Scale bar 7 µm. **(c)** Distribution of nuclei center of mass along the apical-basal axis. **(d)** Nuclear ellipticity and volume distributions along the apical-basal axis. **(e)** Average cross-sectional area distribution along the apical-basal axis for all cells (black) and nuclei (blue). **(f)**. Nuclear and cellular volumes of 140 segmented cells are correlated (r=0.79). **(g)**. The cell and corresponding nuclear cross-sectional areas along the apical-basal axis are highly correlated (r=0.94). **(h)**. An increase in the nuclear cross-sectional area increases the frequency of a neighbour-number-increasing spatial T1 transitions, and vice versa. **(i)** The largest number of changes in neighbour relationships occur at the apical and basal limits of the nucleus for all cells, where cross-sectional areas change sharply. the quadratic law (yellow line).

We conclude that the positions of nuclei can explain much of the observed variability in the cross-sectional cell areas. During the cell cycle, nuclei migrate, and the cell volumes first increase, and subsequently halve due to cell division. As all these processes affect the cross-sectional areas of the cells along the apical-basal axis, one would expect continuous spatial-temporal T1L transitions in growing pseudostratified epithelia.

### 3D cell organisation in growing epithelia

To follow 3D cellular dynamics during epithelial growth and deformation, we cultured embryonic lungs from a Shh^GC/+^; ROSA^mT/mG^ background and imaged every 20 minutes for a total of 10 hours using light-sheet microscopy (Video 6). We used a subset of this dataset (11 time points, >3 hours) (Video 7) to 2.5D segment the apical and basal surfaces, and to explore 3D cell shape dynamics and neighbour relationships in a growing lung bud (Figure 6a, Video 8).

**Figure 6.**
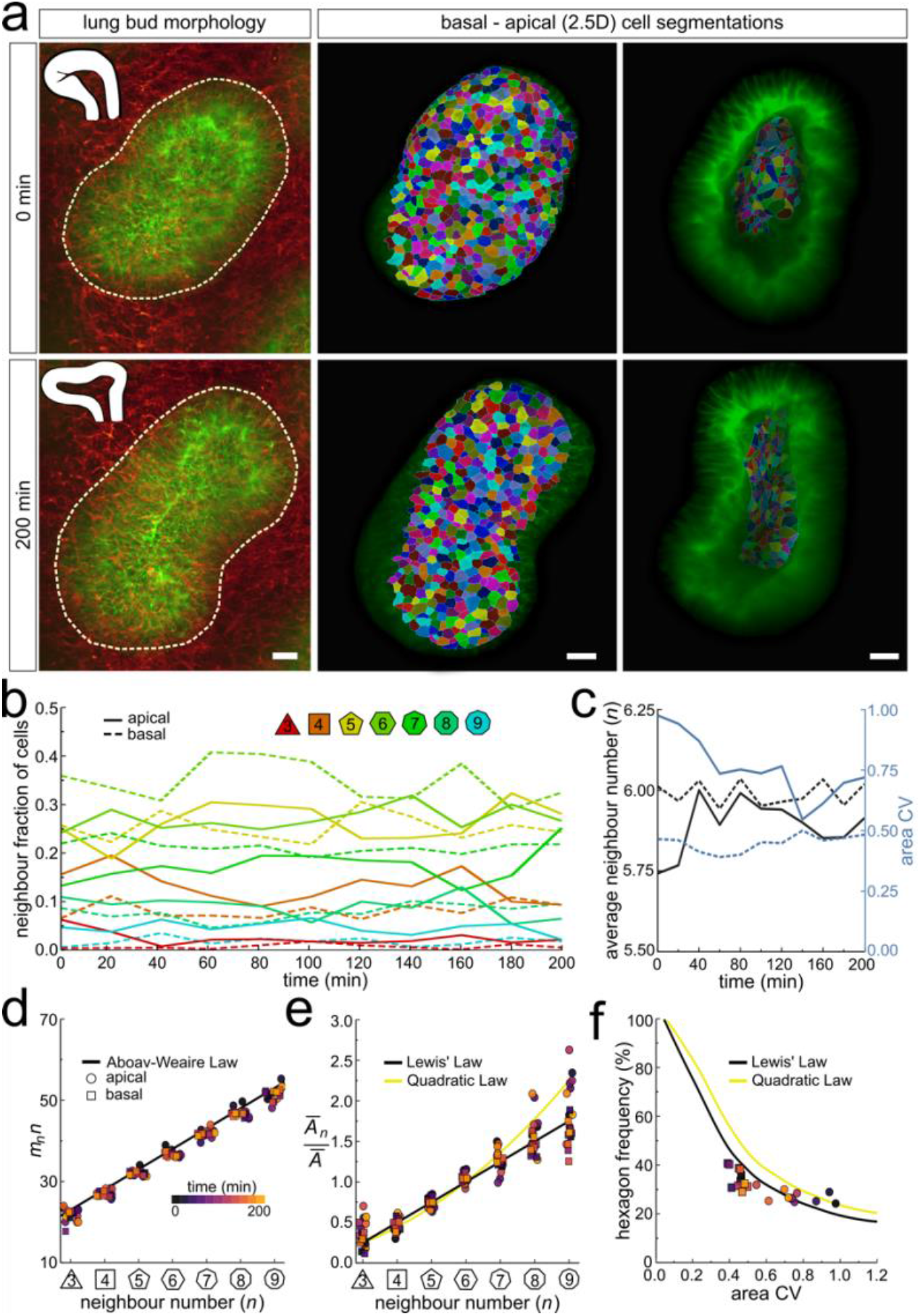
Dynamics of apical and basal epithelial organization. **a)** Timelapse light-sheet microscopy series of a cultured mouse E12.5 distal lung bud expressing the Shh^GC/+^; ROSA^mT/mG^ reporter (green epithelium, and red mesenchyme), imaged every 20 minutes (11 time steps). The white inset denotes the morphology of the lung bud, while the dotted area denotes the segmented cell patch. Corresponding visual provided in Video 6. Cells on both the apical and basal domains were 2.5D segmented, and their morphology quantified. Corresponding visual provided in Video 8. Scale bars 20 µm.(b)Cell neighbour frequencies for the apical and basal layers over time. **(c)** Observed average neighbour number and area coefficient of variation (CV) for the apical and basal layers over time. **(d)** Growing apical and basal layers follow the AW law (black line). Colour code applies to e-f. **(e)** The relative average apical and basal cell areas are linearly related to the number of neighbours (in all time points) and follow Lewis’ law (black line), or the quadratic relationship in the case of higher area variability (yellow line). **(f)** Observed fraction of hexagons versus area coefficient of variation (CV) on the apical and basal layers. The lines mark the theoretical prediction if polygonal cell layers follow either the linear Lewis’ law (black line) or

As the explant was growing, we readjusted the 2.5D segmented region such that the segmented surface area and cell numbers remained roughly constant over time (Figure 6 -figure supplement 1a). Nonetheless, the segmented bud increased in volume as the thickness of the layer increased with time (Figure 6 -figure supplement 1b). Much as in the static dataset, the neighbour number distributions (Figure 6b), and variability of cross-sectional areas (Figure 6c) differ between the apical and basal cell layers in all time points. However, for all time points, both the apical and basal layers conformed to Euler’s formula (Figure 6c), Aboav-Weaire’s law (Figure 6d), and Lewis’ law (Figure 6e). Furthermore, the fraction of hexagons also followed from the variability of the cross-sectional areas, as predicted by the theory (Figure 6f).

We next sought to analyse the 3D dynamics of segmented epithelial cells. As the tracking of packed cells in growing pseudostratified epithelia is challenging, we focused on a small patch with 15 cells in total (Figure 7a). Sequential contour surfaces were drawn to follow cell membrane outlines on several planes along the apical-basal axis and interpolated to reconstruct 3D morphology for each time point (Figure 7 -figure supplement 1, Video 9). All planar segmentations were then pooled into 5 groups along the apical-basal axis to enable morphometric analysis in different tissue regions. Over the time course, the volume of individual cells varied between roughly 400 and 800 µm^3^ (Figure 7b), and the apical-basal length varied between roughly 20 and 30 µm (Figure 7c). We note that all layers conform to Aboav-Weaire’s law (Figure 7d), and Lewis’ law (Figure 7e) in all time points. Moreover, consistent with our theory, the fraction of hexagons follows from the variability of the cross-sectional area, though more deviations are observed, given the small number of cells analysed (Figure 7f).

**Figure 7.**
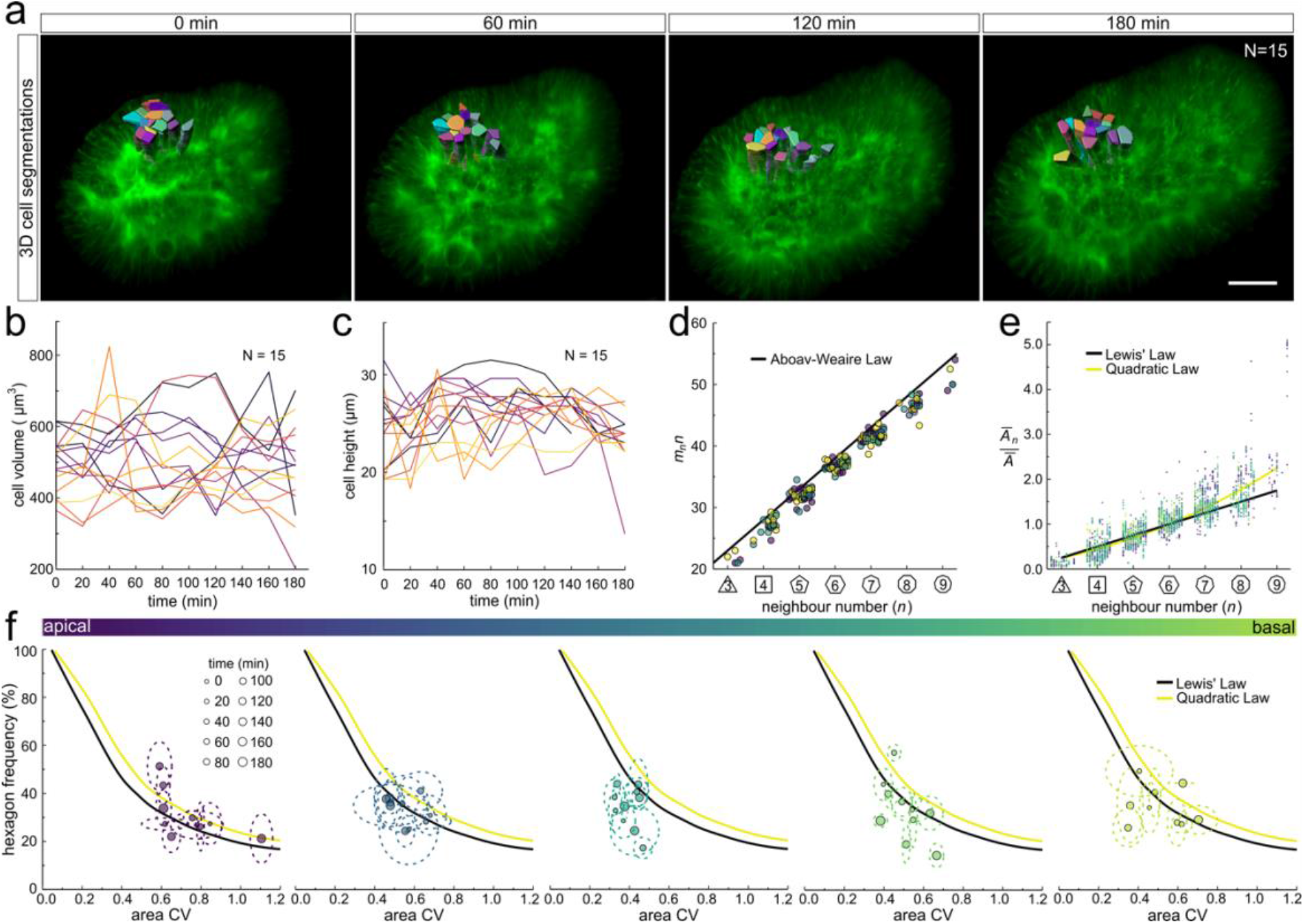
3D cell organization in growing epithelia. **(a)** 3D segmentation of 15 epithelial cells from timelapse light-sheet microscopy imaging of a mouse E12.5 distal lung bud expressing the Shh^GC/+^; ROSA^mT/mG^ reporter. The specimen was imaged every 20 minutes over 3 hours. Planar segmentations along the apical-basal axis were pooled into 5 groups to enable morphometric analysis in different tissue regions. A full timelapse panel is provided in Figure 7 -figure supplement 1 and Video 9. Scale bar 30 µm. **(b)** Epithelial cell volume, and **(c)** height over time (N=15). **(d)** Segmented cells in pooled layers along the apical-basal axis follow the AW law (black line) over time (left to right); see panel f for colour code. **(e)** The relative average cell area in each layer is linearly related to the number of neighbours for all time points (left to right) and follows Lewis’ law (black line), or the quadratic relationship in the case of higher area variability (yellow line); see panel f for colour code. **(f)** Temporal dynamics of observed fraction of hexagons versus area coefficient of variation (CV) along the apical-basal axis. Dotted lines denote variation per time point. Solid lines mark the theoretical prediction if polygonal cell layers follow either the linear Lewis’ law (black line) or the quadratic law (yellow line).

Much as in the static dataset (Figure 3,4), we observe up to 14 neighbour number changes (T1L transitions) along the apical-basal axis (Figure 8a,b). The average number of T1L transitions is relatively constant over time (Figure 8b). The mean relative apical-basal position for T1L transitions is again roughly in the middle, but in this small dataset, we now observe more T1L transitions in the center of the cell than at the apical or basal boundaries (Figure 8c). By following a single cell over time, we can appreciate the dynamic cell shape changes, and how a change in the cross-sectional area correlates with a change in neighbour number (Figure 8d). The neighbour relationships are, of course, not determined by the local cell cross-section, but by the overall cross-sectional area distribution in that layer. Accordingly, the correlation between the cross-sectional area and the neighbour number is not perfect for a single cell. By considering a patch of cells, we can, however, see how those T1L transitions occur dynamically in developing tissues (Figure 8e).

**Figure 8.**
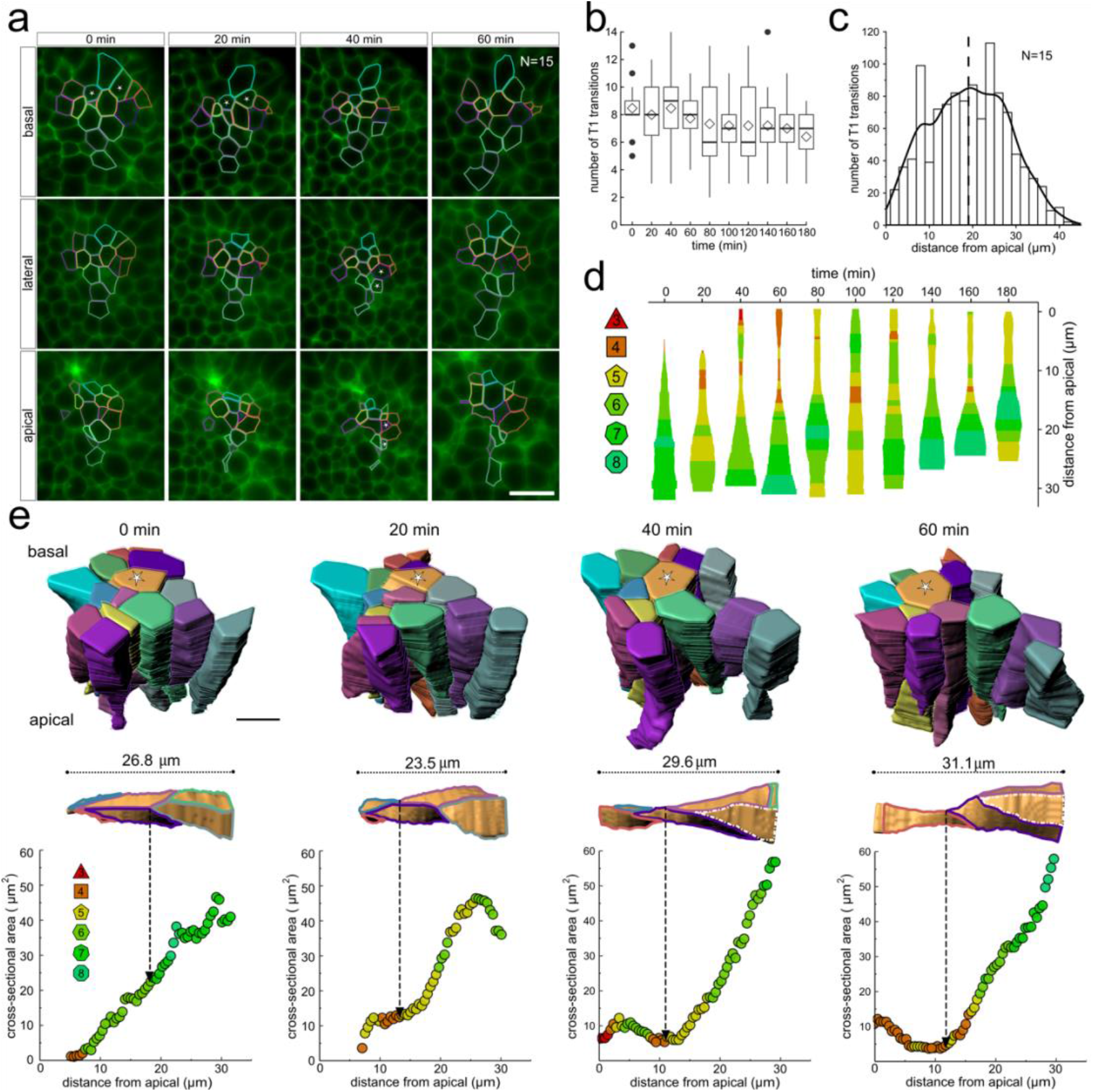
3D cell neighbour dynamics in growing epithelia. **(a)** Apical, basal and lateral cross-sections from a light-sheet microscopy timelapse of a murine E12.5 distal lung bud expressing the Shh^GC/+^; ROSA^mT/mG^ reporter. The specimen was imaged every 20 minutes over 3 hours. Cell membrane outlines illustrate fluid cell neighbour relationships along the apical-basal axis and over time. T1 transitions are marked with white stars; see panel e for cell colour code. Scale bar 14 µm. **(b)** Number of T1L transitions for all cells (N=15) over time. Diamonds represent the mean. Morphometric quantifications of planar segmentations along the apical-basal axis were used to examine T1L transition dynamics. **(c)** Spatial distribution along the apical-basal axis of T1L transitions for all cells and time points. **(d)** Temporal evolution of neighbour relationships along the apical-basal axis for a single cell. Schematic cell width corresponds to cross-sectional area. **(e)** (top row) 3D iso-surface segmentations of 15 epithelial cells. Scale bar 10 µm. (bottom row) Cross-sectional area and cell neighbour number along the apical-basal axis for a given cell (marked with a white star). Dotted lines indicate contact with a cell that was not segmented.

## DISCUSSION

Epithelial tissues remodel into complex geometries during morphogenesis. We used light-sheet microscopy and 3D cell segmentation to unravel the physical principles that define the 3D cell neighbour relationships in pseudostratified epithelial tissues. Our analysis reveals that pseudostratified epithelial layers adopt a far more complex packing solution than previously anticipated: the 3D epithelial cell shapes are highly irregular, and cell neighbour relationships change multiple times along the apical-basal axis, with some cells having up to 14 changes in their neighbor contacts along their apical-basal axis (Figure 3a). Curvature effects can result in neighbour changes, but the data does not show the dependency on cell neighbour numbers that would be expected if curvature effects played a dominating role (Figure 3g). There is also no apical-basal bias (Figure 3e), and the prevalence of contact remodeling is randomly distributed (Figure 4i).

Even though the neighbour relationships are uncorrelated between the apical and basal sides and appear random at first sight, they follow the same fundamental relationships that have previously been described for apical epithelial layers, i.e. Euler’s formula, Lewis’ law, and Aboav-Weaire’s law across the entire tissue and at all times (Figure 2, 4, 6, 7). This arrangement minimizes the lateral cell-cell surface energy in each plane along the apical-basal axis, given the variability in the cell cross-sectional areas (Figure 1) (Kokic et al., 2019; Vetter et al., 2019). Where present, the stiff nucleus determines the cell cross-sectional area, as is apparent from the strong correlation between the cell cross-sectional and the nuclear cross-sectional areas (Figure 5g). Accordingly, most changes in neighbour relationships occur at the apical and basal limits of the nucleus where cross-sectional areas change sharply (Figure 5i). As the nucleus moves along the apical-basal axis during the cell cycle, a process referred to as interkinetic nuclear migration (IKNM) (Meyer et al., 2011), cell neighbour relationships change continuously (Figure 8). We conclude that neighbour relationships in epithelia are fluidic, and the complex, dynamic 3D organisation of cells in growing epithelia follows simple physical principles.

Defining the physical principles behind cell neighbour relationships is only the first step in unravelling the determinants of epithelial 3D cell shapes. The second key aspect is the cell volume distribution along the apical-basal axis, which gives rise to the cell cross-sectional area distribution, which then determines the cell neighbour relationships (Figure 5e). The overall cell volume is determined by cell growth and division, but its distribution along the apical-basal axis depends on the nuclear dynamics (Figure 5g), and the epithelial cell heights. We find that the nucleus occupies, on average, 55% of the cell volume in the embryonic lung epithelia. As the cell nuclei move along the apical-basal axis during the cell cycle (Meyer et al., 2011), the cytoplasm fills the remaining space between the apical and basal surfaces, likely in a way that minimises the total surface area of all cells. The determinants of the epithelial thickness, i.e. the distance between the apical and basal surfaces are still unknown, but signalling factors that control cellular tension are known to affect cell height (Widmann and Dahmann, 2009).

Cell-based modelling frameworks are heavily used to investigate epithelial processes and how they result in morphological changes such as tissue bending, folding, fusion, and anisotropic growth during morphogenesis (Fletcher et al., 2014; Tanaka, 2015). Our data confirms many underlying assumptions of cell-based modelling frameworks and provides quantitative data to calibrate parameters. Once calibrated to reproduce the here identified 3D cell shape distributions, such simulation frameworks will help to reveal the determinants of 3D cell shapes, and will be invaluable in providing insight into how local changes in cell growth, adhesion, tension, or in the basal lamina affect cell shapes locally and within the remaining epithelial layer.

In summary, this study offers a detailed view of 3D cell neighbor relationship dynamics and packing in growing epithelial tissues, and demonstrates that the 3D cell shapes are much more complex than previously anticipated, and that cell neighbor relationships are dynamic and change as result of cell growth and cell cycle-linked IKNM. The complex 3D cell neighbor relationships can nonetheless be understood based on simple physical principles. Although we recognize that tissue architecture is a multifactorial process, our work carries vast implications for the study of cell-cell signaling, epithelial cohesion, and energetic modeling of developing epithelial layers in both healthy and disease contexts.

## MATERIALS AND METHODS

### Ethical Statement

Permission to use animals was obtained from the veterinary office of the Canton Basel-Stadt (license number 2777/26711). Experimental procedures were performed in accordance with the Guide for the Care and Use of Laboratory Animals and approved by the Ethics Committee for Animal Care of ETH Zurich. All animals were housed at the D-BSSE/UniBasel facility under standard water, chow, enrichment, and 12-hrs light/dark cycles.

### Animals

To investigate 3D cellular dynamics during mouse embryonic lung development, we used mouse lung rudiments from animals homozygous for the ROSA^mT/mG^ and heterozygous for the Shh-cre allele (Shh^cre/+^; ROSA^mT/mG^). The double-fluorescent Shh-controlled Cre reporter mouse expresses membrane-targeted tandem dimer Tomato (mT) before Cre-mediated excision and membrane-targeted green fluorescent protein (mG) after excision (Muzumdar et al., 2007). As a result, only epithelial cell membranes are labelled by GFP, while all adjacent mesenchymal tissue is labelled by tdTomato.

### Immunofluorescence

E12.5 mouse lungs were fixated for 1hr in 4% paraformaldehyde in PBS, and subsequently incubated with Lamin B1 (Thermo; Material No. 702972; 1:200) at 4 °C for three days. As a structural component of the nuclear lamina, LaminB1 immunostaining makes crowded nuclei clearly distinguishable and easily segmentable. After washing in D-PBS, lungs were incubated with conjugated fluorescent secondary Alexa Fluor 555 donkey anti-mouse IgG (H+L) (Abcam; Material No. ab150106; 1:250) for two days at 4 °C.

### Optical clearing and Lightsheet imaging

Optical clearing of embryonic lung rudiments enabled the 3D segmentation of numerous epithelial cells from single image stacks. To this extent, the whole-mount clearing of dissected E12.5 lung explants was performed with the Clear Unobstructed Brain/Body Imaging Cocktails and Computational Analysis (CUBIC) protocol (Susaki et al., 2015) (Figure 2 -figure supplement 3). Reagents for delipidation and refractive index (RI) matching were made as follows: CUBIC-1 [25% (w/w) urea, 25% ethylenediamine, 15% (w/w) Triton X-100 in distilled water], and CUBIC-2 [25% (w/w) urea, 50% (w/w) sucrose, 10% (w/w) nitrilotriethanol in distilled water], respectively. Following fixation and immunostaining, samples were incubated in 1/2 CUBIC-1 (CUBIC-1:H2O=1:1) for four days, and in 1X CUBIC-1 until they became transparent. All explants were subsequently washed several times in PBS and treated with 1/2 CUBIC-2 (CUBIC-2:PBS=1:1) for around four days. Lastly, incubation in 1X CUBIC-2 was done until the desired transparency was achieved. All solutions were changed daily, and CUBIC-1 steps were performed on a shaker at 37 °C while CUBIC-2 steps at room temperature. Cleared samples were then embedded in 2% low melting point solid agarose cylinders and immersed in CUBIC-2 for two more days to increase the agarose refractive index. 3D image stacks were acquired on a Zeiss Lightsheet Z.1 microscope using a Zeiss 20x/1.0 clearing objective (Supplementary Figure 1).

### Timelapse light-sheet acquisitions

Light-sheet acquisitions of live epithelial cell morphology enabled the study of 3D organization dynamics. Following dissection in DPBS at room temperature, E12.5 lung explants were cultured in sterile Dulbecco’s modified Eagle’s medium w/o phenol red (DMEM) (Life Technologies Europe BV; 11039021) containing 10% Fetal Bovine Serum (FBS) (Sigma-Aldrich Chemie GmbH; F9665-500ML), 1% Glutamax (Life Technologies Europe BV; A1286001), and 1% penicillin/streptomycin (Life Technologies Europe BV; 10378-016). All specimens were equilibrated at 37°C with 5% CO2 in a humidified incubator for 1hr.

Following a 1hr equilibration period, 1.5% LMP hollow agarose cylinders were prepared (Udan et al., 2014). Hollow cylinders, in contrast to solid ones, accommodate unencumbered 3D embryonic growth, provide boundaries to minimize tissue drift, enable imaging from multiple orientations, and allow for better perfusion of gasses and nutrients. All specimens were suspended within each hollow cylinder in undiluted Matrigel (VWR International GmbH; 734-1101), an ECM‐based optically clear hydrogel that provided a near-native 3D environment and supported cell growth and survival. All cylinders were kept at 37°C with 5% CO2 in culture media for 1hr before mounting.

For each overnight culture, the imaging chamber was prepared by sonication at 80°C, followed by ethanol and sterile PBS washes. After assembly, the chamber was filled with culture medium and allowed to equilibrate at 37°C with 5% CO2 for at least 2hrs before a cylinder containing an explant was mounted for imaging. Furthermore, to compensate for evaporation and to maintain a fresh culture media environment, two peristaltic pumps were installed to supply 0.4 mL and extract 0.2 mL of culture medium per hour. Each lung explant was then aligned with the focal plane within the center of a thin light-sheet to enable fine optical sectioning with optimal lateral resolution. For this study, all live imaging was done with a 20x/1.0 Plan-APO water immersion objective.

### Image processing

To efficiently process the resulting volumetric CZI datasets (10s-100s of GBs), all image stacks were transferred to a storage server and subsequently processed in remote workstations (Intel Xeon CPU E5-2650 with 512 GB memory). Deconvolution via Huygens Professional v19.04 (Scientific Volume Imaging, The Netherlands, http://svi.nl) improved overall contrast and resolution while Fiji (ImageJ v1.52t) (Schindelin et al., 2012) aided in accentuating cell membranes, enhancing local contrast, removing background fluorescence, and TIFF conversion.

### Cell morphometric quantifications

Cell morphology on the apical and basal membranes of embryonic lung epithelia was investigated using the open-source software platform MorphoGraphX (MGX) (Barbier de Reuille et al., 2015). By meshing the curved boundaries of input 3D image stacks and projecting nearby signal onto it, MGX builds a curved 2.5D image projection that is distortion-free, unlike planar 2D projections that ignore curvature. We then proceeded to use a suitable implementation of the Watershed transform to extract individual cell geometries, with minimal manual curation, and quantify properties such as surface area and the number of cell neighbours. All border cells were excluded. Apical and basal cell meshes were exported as text files and traversed using the R Programming Environment to extract the neighbour relationships between cells as needed to generate Aboav-Weaire plots.

To render time-lapse datasets and extract 3D volumetric surface reconstructions of entire epithelial cells, we employed Imaris v9.1.2 (Bitplane, South Windsor, CT, USA). By computationally interpolating between cell membrane contour surfaces from successive transverse frames into iso-surfaces, faithful cell and nuclear 3D volumes were obtained. Quantified volumetric features included cell and nuclear volume, total surface area, sphericity, and nuclear position along the apical-basal axis. Imaris was also used to generate high-resolution videos, which, despite being strongly downsampled to accommodate vast time-lapse datasets, presented little noticeable loss in image quality. Furthermore, to extract cell areas and the number of neighbours along the apical-basal axis, transverse image frames were imported into ImageJ and processed using the interactive plugin TissueAnalyzer (Aigouy et al., 2016). Like this, cell segmentation masks across layers could be generated, and cell geometry and neighbour topology quantified.

## Acknowledgements

This work has been supported through an SNF Sinergia grant to DI. We acknowledge the many discussions and varied input of Marco Kokic, Anđela Markovic, Odyssé Michos, and Erand Smakaj, and we thank Richard Smith for help with MorphographX.

## Author Contributions

DI and RV developed the theoretical framework. HG obtained the cleared light-sheet data, MD obtained the live imaging data with support from HG. HG, MD, and LH processed and analysed the data. DI wrote the manuscript with contributions from RV; HG wrote the Methods section and figure legends; all authors approved the final manuscript.

## FIGURE SUPPLEMENTS

**Figure 2 - figure supplement 1.**
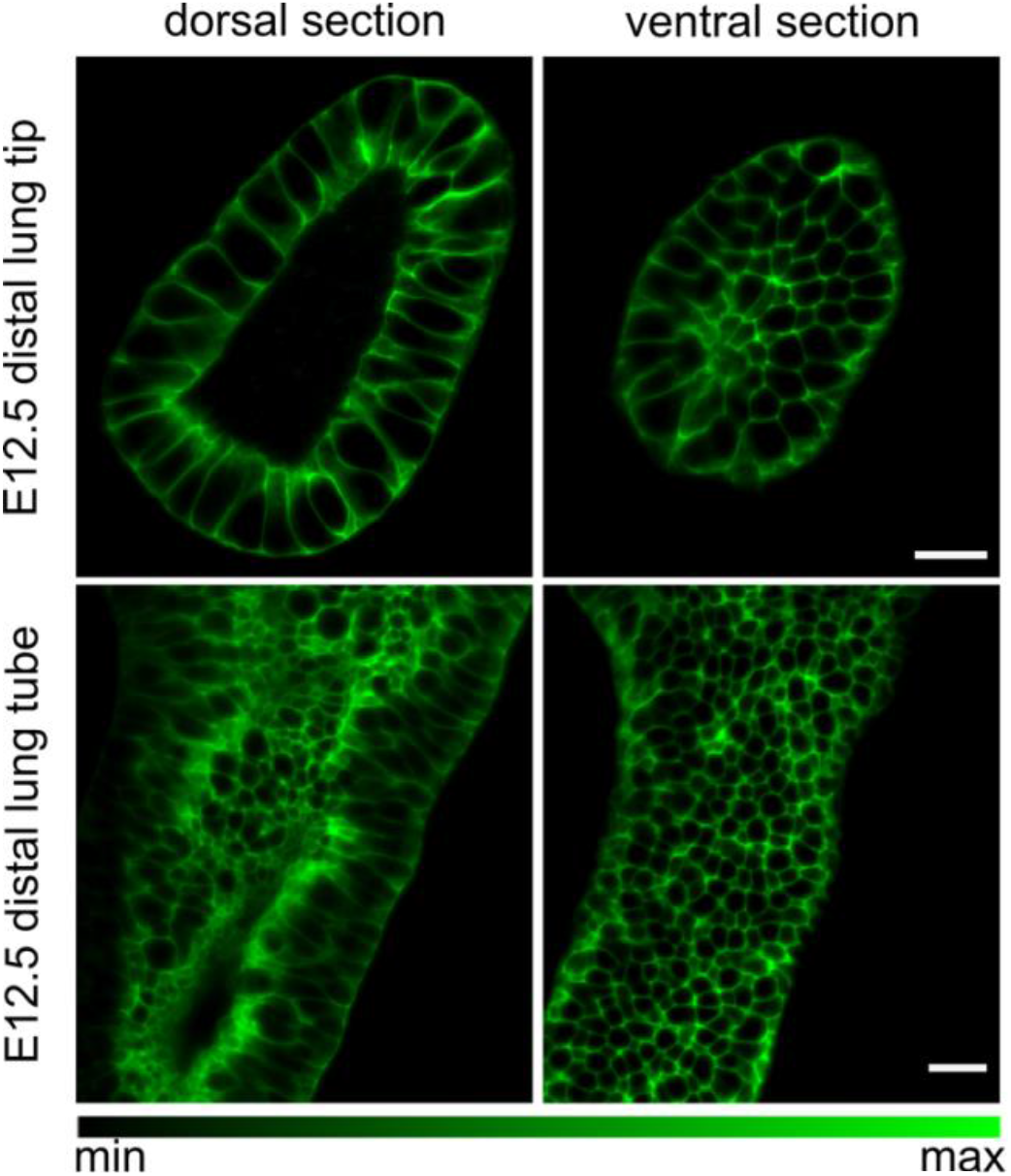
Embryonic mouse lung rudiments. . Dorsal and ventral cross-sections of an E12.5 lung tip and tube carrying the Shh^GC/+^; ROSA^mT/mG^ reporter allele, marking the epithelial lineage. Tissue explants were optically cleared using CUBIC and imaged using light-sheet microscopy to achieve cellular resolution at all depths. Scale bars 20 µm.

**Figure 2 - figure supplement 2.**
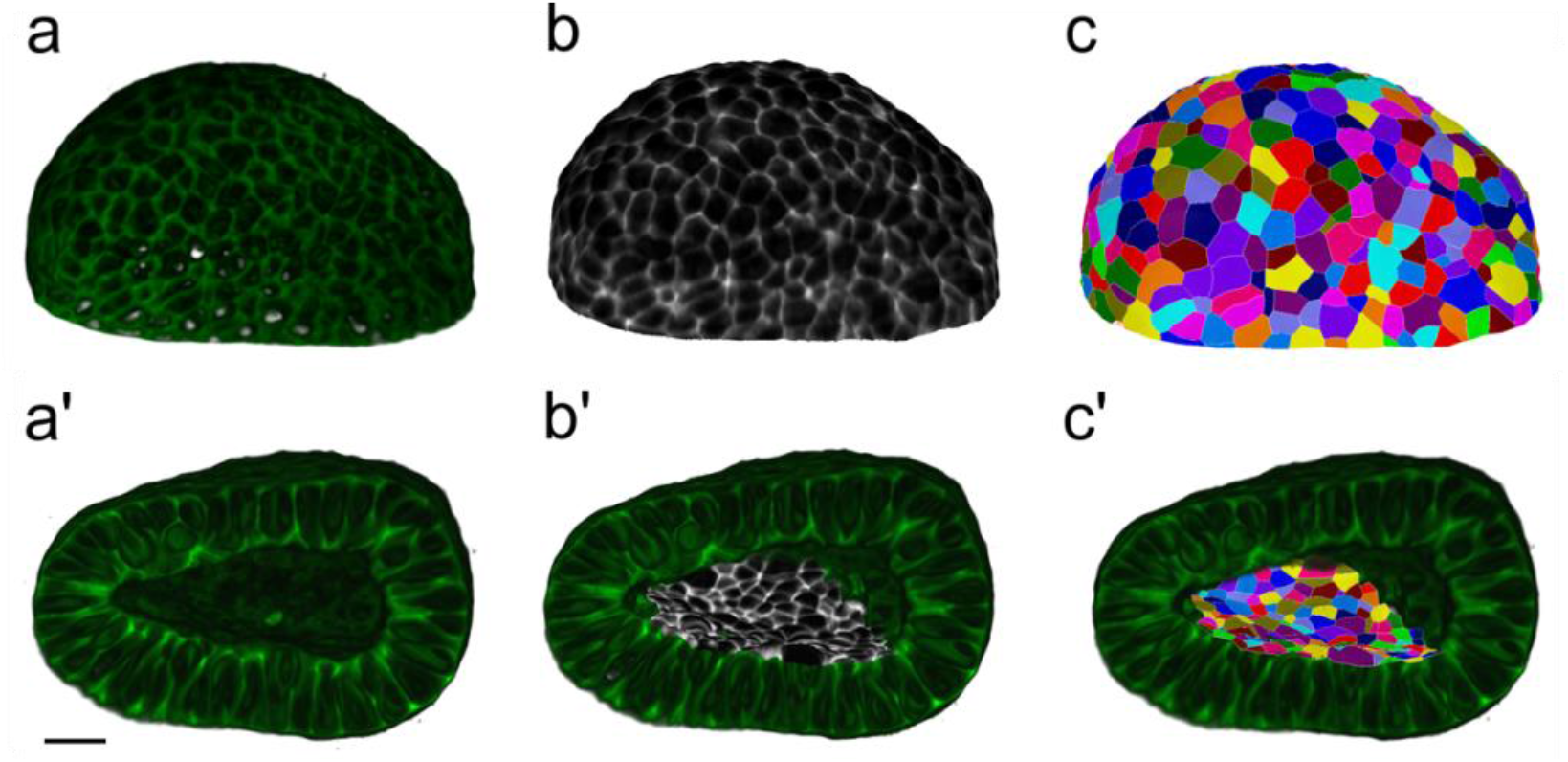
Workflow for surface cell segmentations. . MorphoGraphX segmentation workflow to extract cell segmentations along curved surface boundaries (2.5D) for a CUBIC-cleared murine E12.5 distal lung tip. The illustrated sample expressed the Shh^GC/+^; ROSA^mT/mG^ reporter and was imaged using light-sheet microscopy. **(a-c)** Correspond to the basal layer while **(a’-c’)** correspond to the apical domain. **(a, a’)** Deconvolved light-sheet microscopy 3D renderings showing the basal and apical surfaces. **(b, b’)** Curved surfaces are isolated, meshed, and the adjacent fluorescent signal is projected. **(c, c’)** Apical and basal domains are segmented (2.5D segmentations) and colored by random label numbers. Scale bar 20 µm.

**Figure 2 - figure supplement 3.**
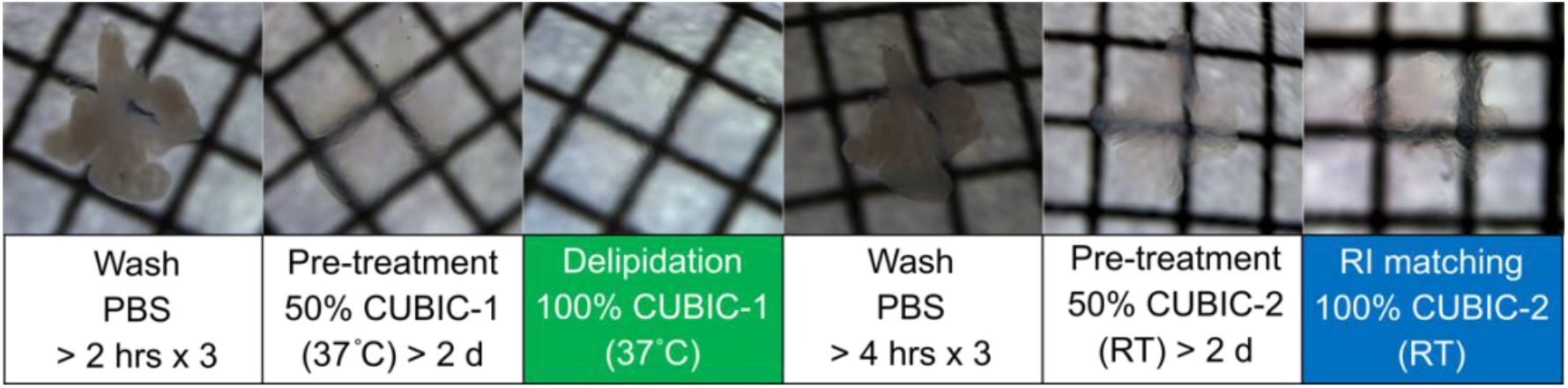
CUBIC clearing of embryonic tissue. . Protocol for advanced CUBIC (Clear, Unobstructed Brain/Body Imaging Cocktails and Computational analysis) of a murine lung rudiment [Susaki _CUBIC_2015]. Serial dilutions in reagent-1 and reagent-2 guarantee that morphology is not distorted and that optical transparency is achieved within one week with high reproducibility.

**Figure 3 - figure supplement 1.**
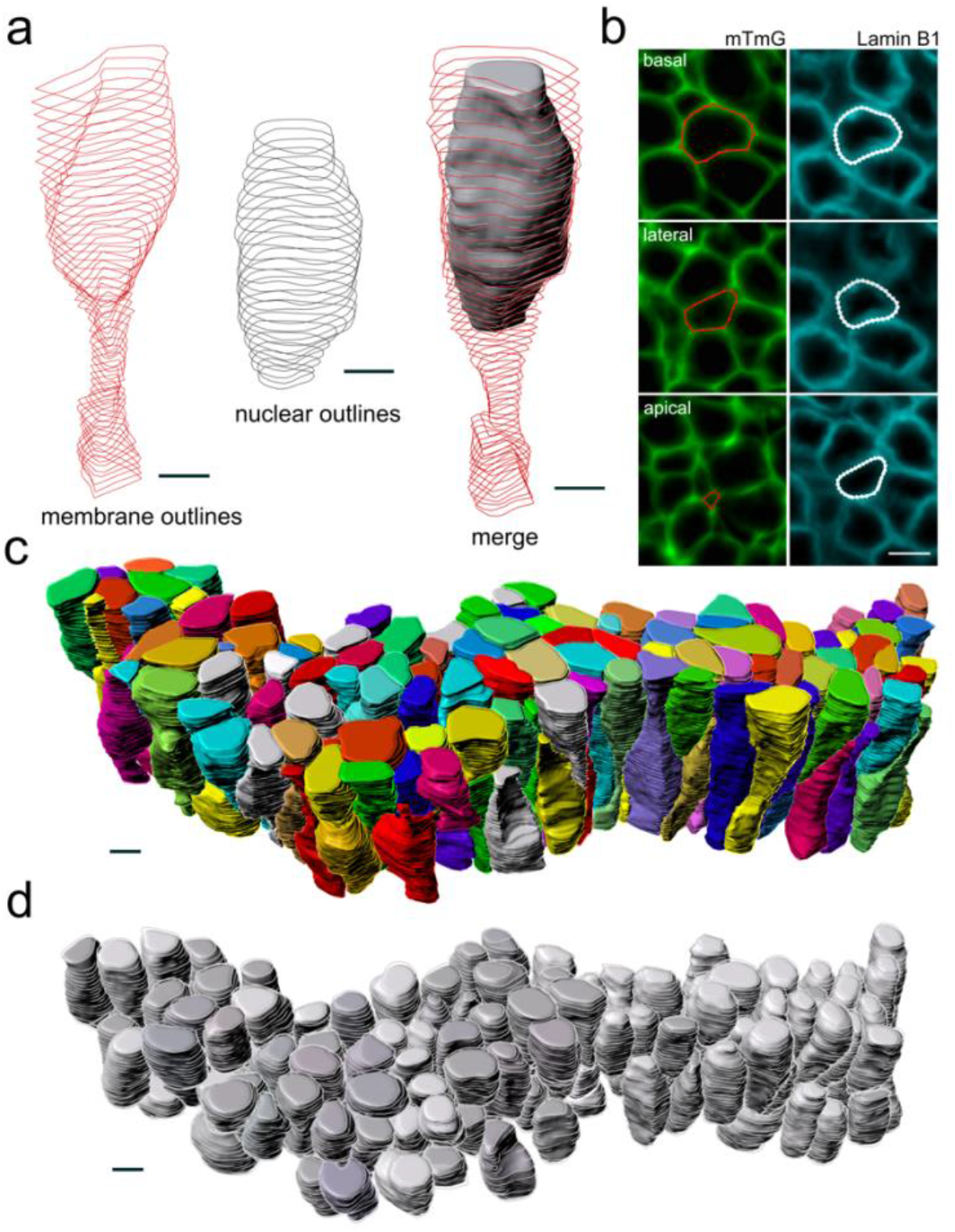
Workflow for 3D epithelial cell and nuclear segmentations. . Segmentation workflow to extract cellular and nuclear 3D shapes from a CUBIC-cleared murine E12.5 lung tube. The specimen used expressed the Shh^GC/+^; ROSA^mT/mG^ reporter and was immunostained for lamin B1 to both selectively label cell membranes and mark nuclear envelopes. Light-sheet microscopy was used. **(a)** Sequential contour surfaces are drawn to follow cell membrane and nuclear outlines on a number of planes along the apical-basal axis. By interpolating between contours, iso-surfaces accurately representing 3D shapes can be extracted. **(b)** Membrane and nuclear contour surface overlays at different tissue depths. **(c)** Extracted 3D cell, and **(d)** nuclear iso-surfaces (N=140) for developing mouse lung. Co-planar contours were used for morphometric quantifications. All scale bars 5 µm.

**Figure 5- figure supplement 1.**
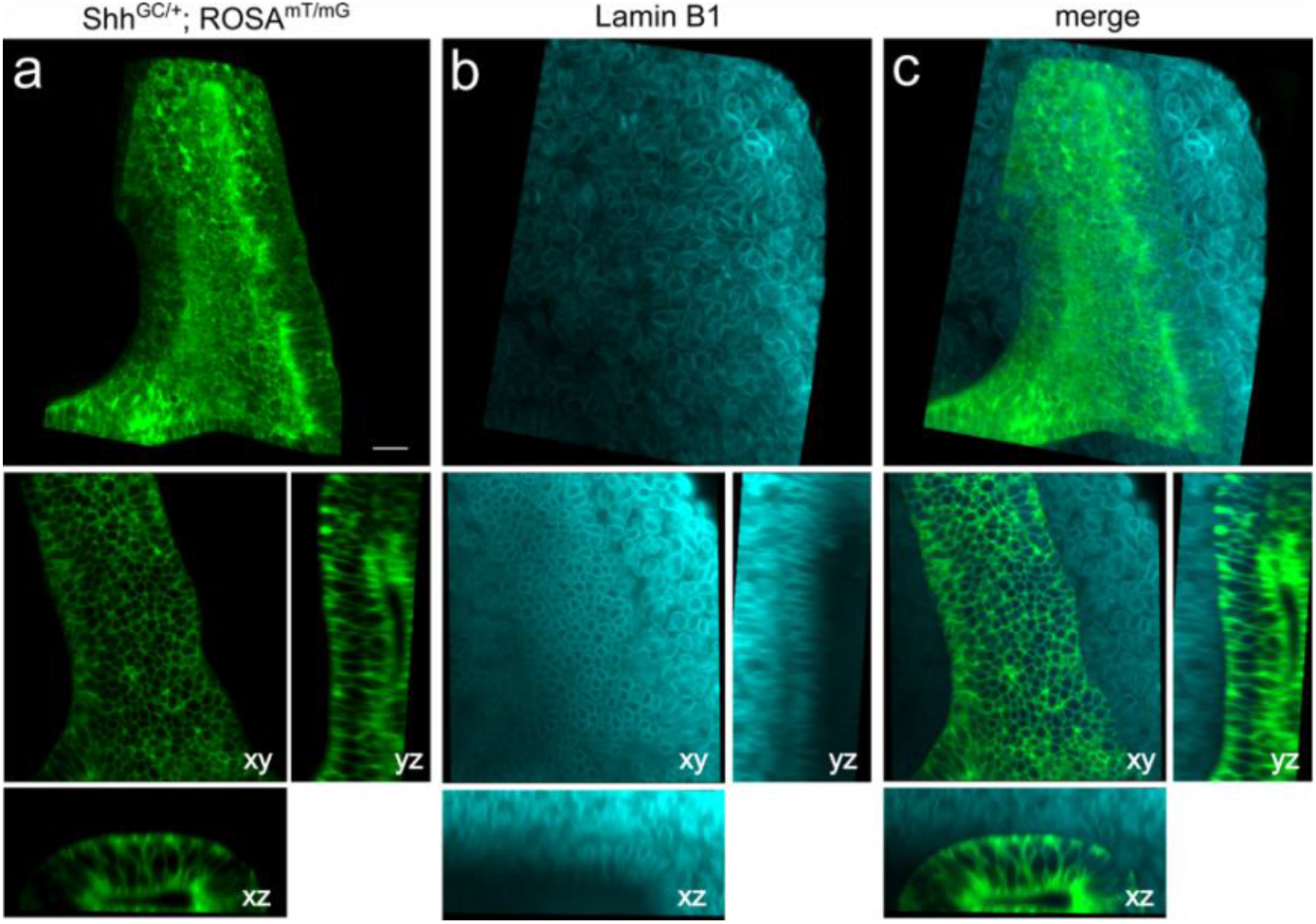
figure supplement 1. Light-sheet imaging of a stained embryonic mouse lung rudiment. . Volumetric renderings and orthogonal projections obtained from light-sheet microscopy for an E12.5 embryonic lung. The above specimen carried the Shh^GC/+^;ROSAm^T/mG^ to mark **(a)** epithelial cell membranes and was **(b)** immunostained for lamin B1 to mark nuclear envelopes. **(c)** merged fluorescent signal. Scale bar 20 µm.

**Figure 6 - figure supplement 1.**
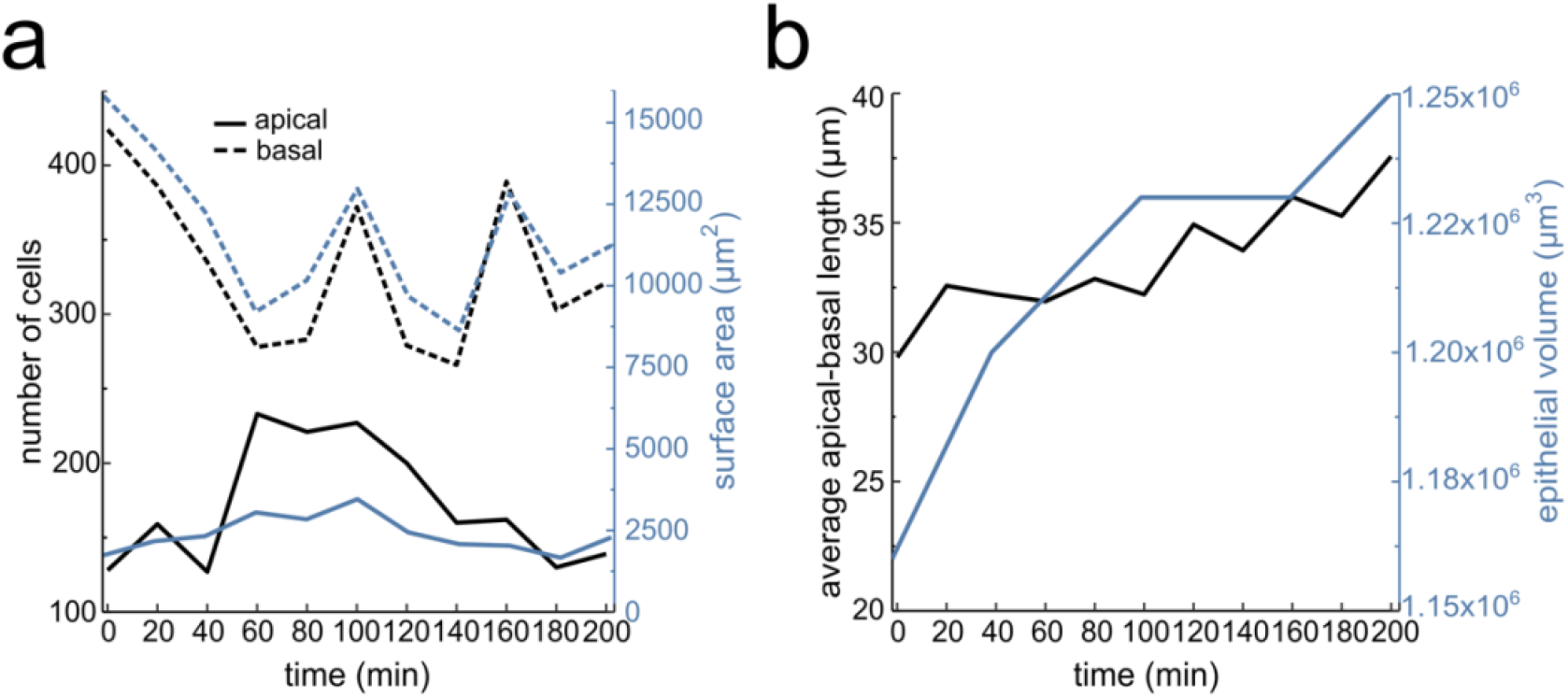
Lung bud viability and growth quantifications over time. **(a)**Number of segmented cells and surface area for the apical and basal layers over time. **(b)** Average apical-basal length and epithelial volume over time. Tissue thickness was measured at five landmark regions along the growing bud.

**Figure 7 - figure supplement 1.**
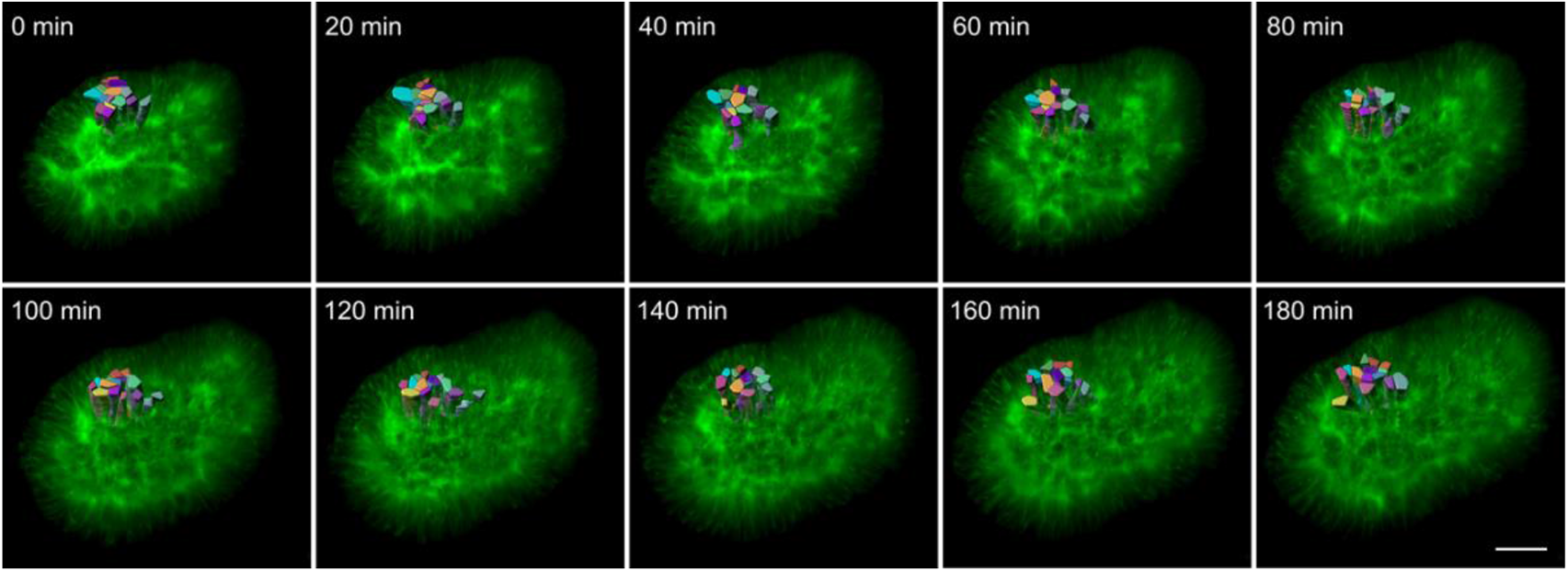
3D timelapse segmentation of growing epithelia. Timelapse segmentation of growing epithelial cells from a CUBIC-cleared mouse E12.5 lung tip over 180 minutes; iso-surfaces colored by cell identifier. The specimen used expressed the Shh^GC/+^; ROSA^mT/mG^ reporter and was imaged using light-sheet microscopy. Scale bar 30 µm.

### VIDEOS

**Video 1.**
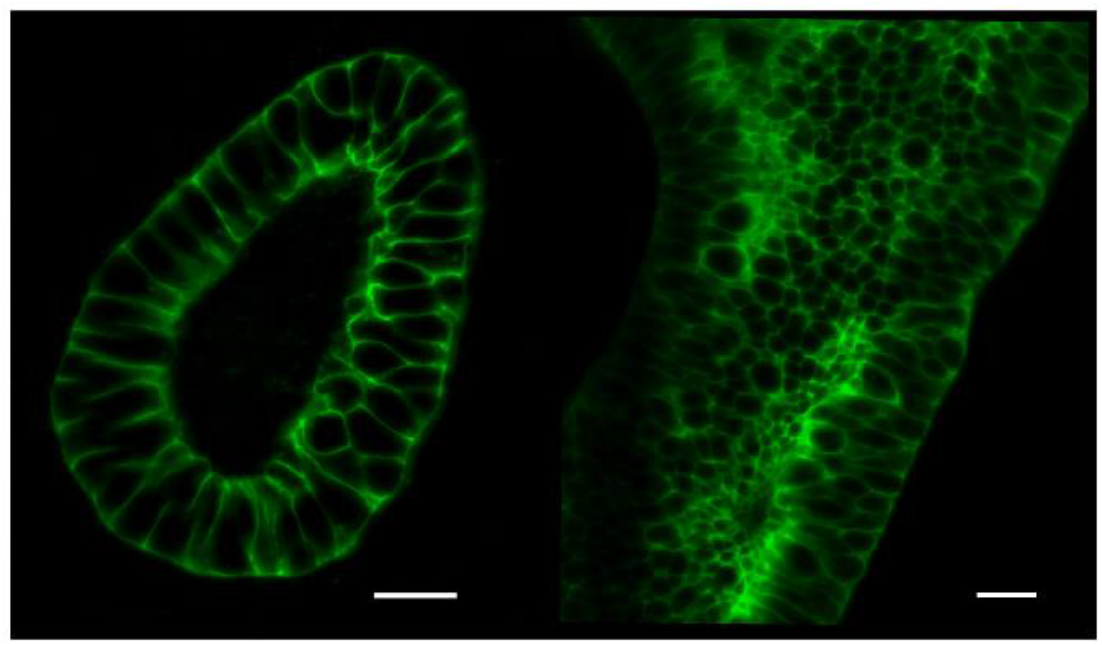
Optically cleared mouse lung rudiment image stacks. Animated cross-section exploration of murine E12.5 lung tip and tube image stacks. Specimens carried the Shh^GC/+^; ROSA^mT/mG^ reporter allele to mark the epithelial lineage were CUBIC cleared, and subsequently imaged using light-sheet microscopy to achieve cellular resolution. Scale bars 20 µm. (AVI 83.1 MB).

**Video 2.**
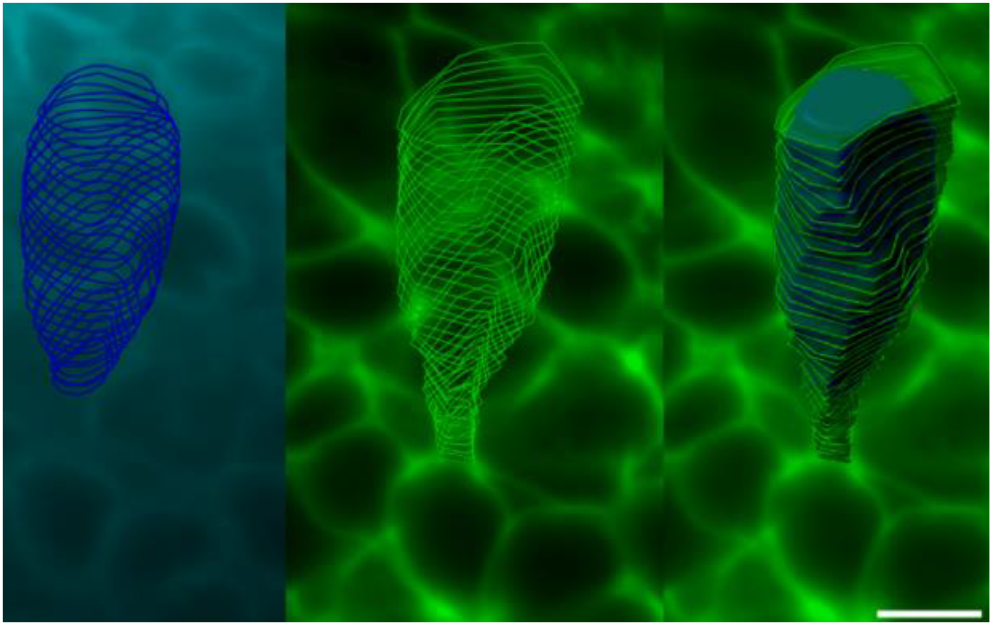
3D contour surfaces. Image stack animation illustrating nuclear and cell contour surface overlays (left, middle) along the apical-basal axis as well as full 3D iso-surfaces (right) for a single cell. A CUBIC-cleared E12.5 Shh^GC/+^; ROSA^mT/mG^ lung tube showing epithelial membranes in green was immunostained for lamin B1 rendering nuclear envelopes blue. Scale bar 5 µm. (AVI 11.8 MB).

**Video 3.**
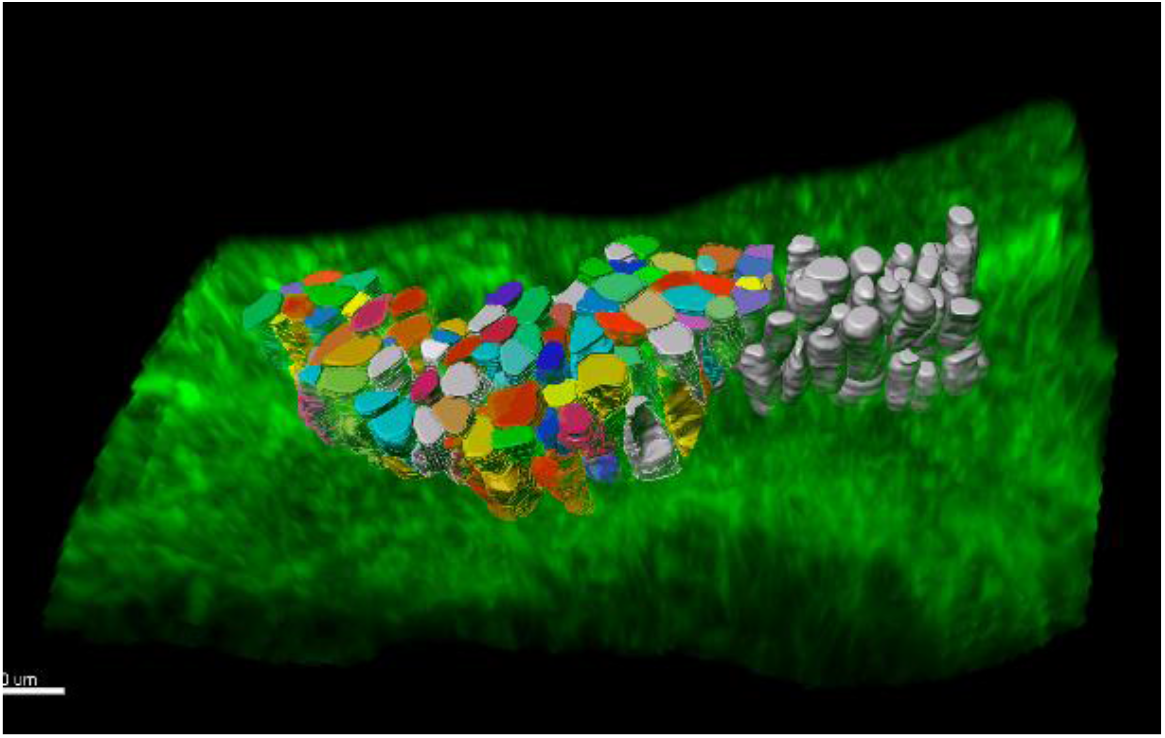
3D nuclear and tube cell segmentations. Volumetric rendering of an E12.5 Shh^GC/+^; ROSA^mT/mG^ lung tube (green epithelia) imaged using light-sheet microscopy. 3D cellular and nuclear segmentations (N=140) were facilitated by CUBIC tissue clearing. (AVI 42.0 MB).

**Video 4.**
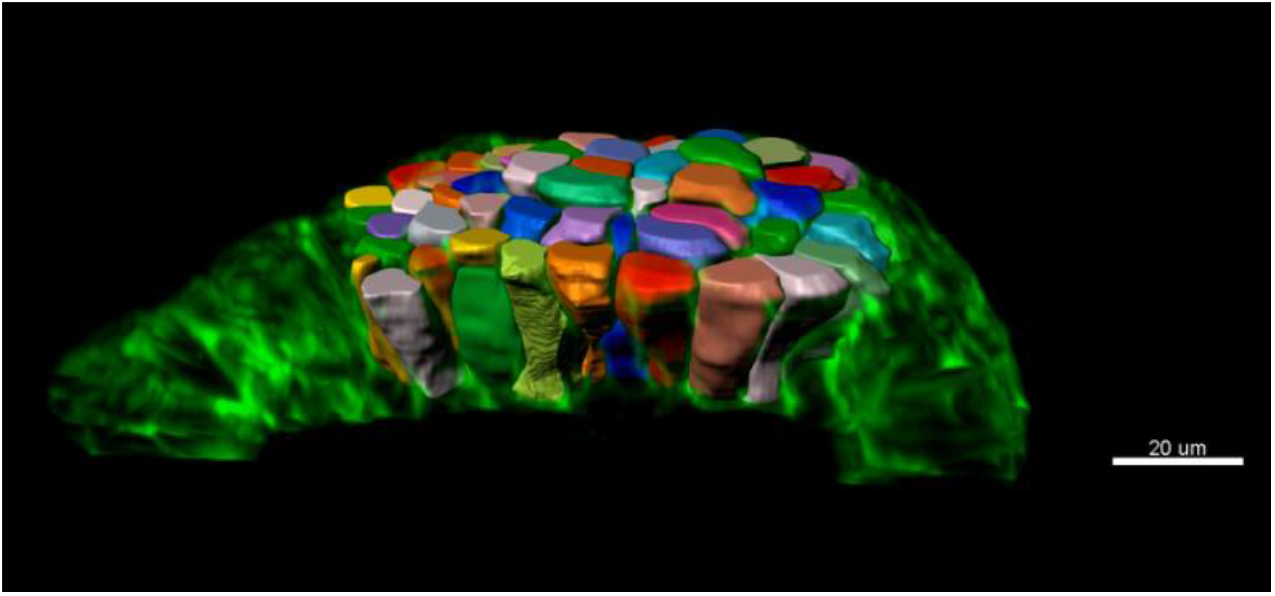
3D tip cell segmentations. Volumetric rendering of an E12.5 Shh^GC/+^; ROSA^mT/mG^ lung tip (green epithelia) imaged using light-sheet microscopy. 3D cellular segmentations (N=59) were facilitated by CUBIC tissue clearing. (AVI 12.4 MB).

**Video 5.**
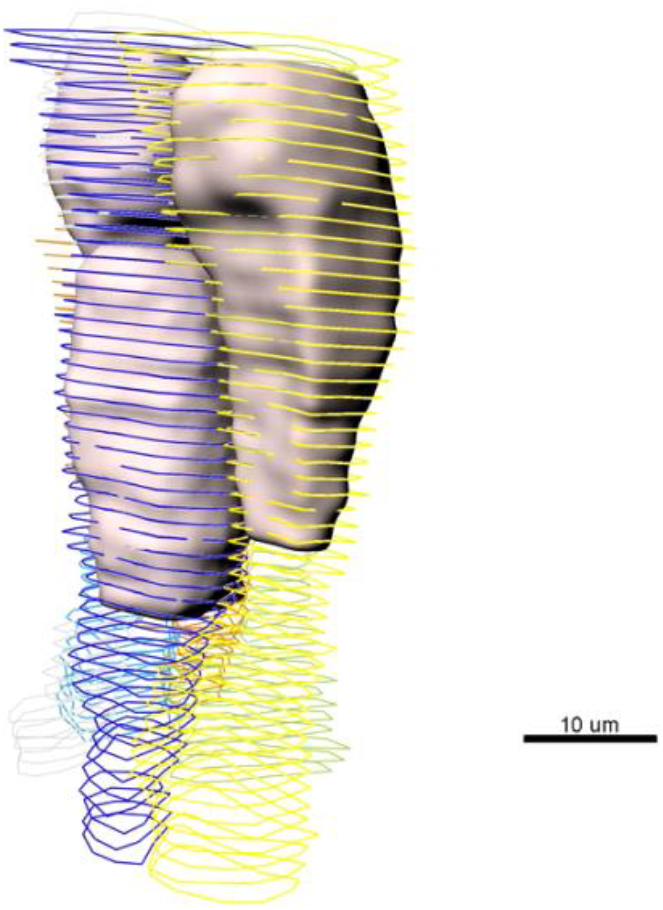
3D nuclear iso-surface segmentations and cell membrane contour. Volumetric rendering of sequential cell membrane contour surfaces and nuclear iso-surfaces from an E12.5 Shh^GC/+^; ROSA^mT/mG^ lung tube immunostained for lamin B1. The sample was cleared using CUBIC and imaged on a Z.1 Zeiss light-sheet microscope. (AVI 8.70 MB).

**Video 6.**
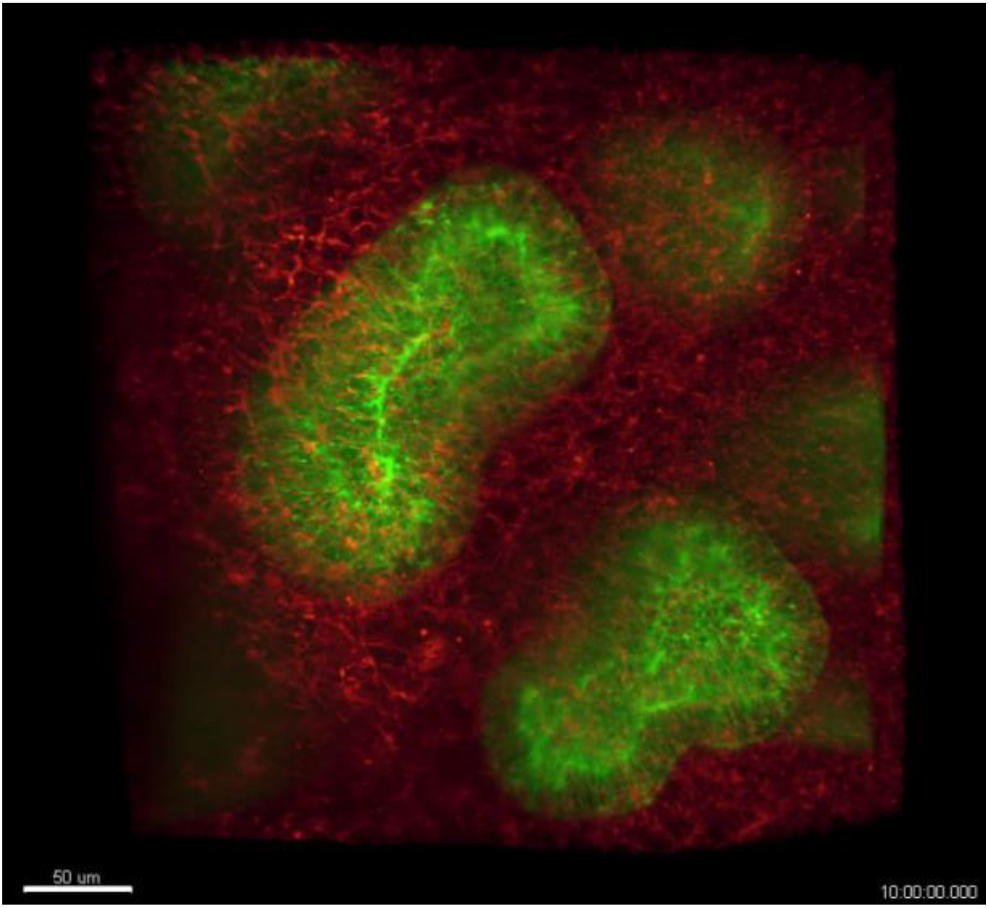
High-resolution light-sheet microscopy timelapse imaging of epithelial lung development (10 hours). Timelapse movie showing the development of an E12.5 mouse lung rudiment carrying the Shh^GC/+^; ROSA^mT/mG^ construct. Embryonic lung was mounted in a hollow cylinder made from low-melting-point agarose and filled with matrigel to replicate the native microenvironment and promote near-physiological growth. Sample was imaged using the Zeiss Z.1 Lightsheet system for 10 hours, with frames acquired every 20 minutes. Top bud was used for quantifications. (AVI 44.5 MB).

**Video 7.**
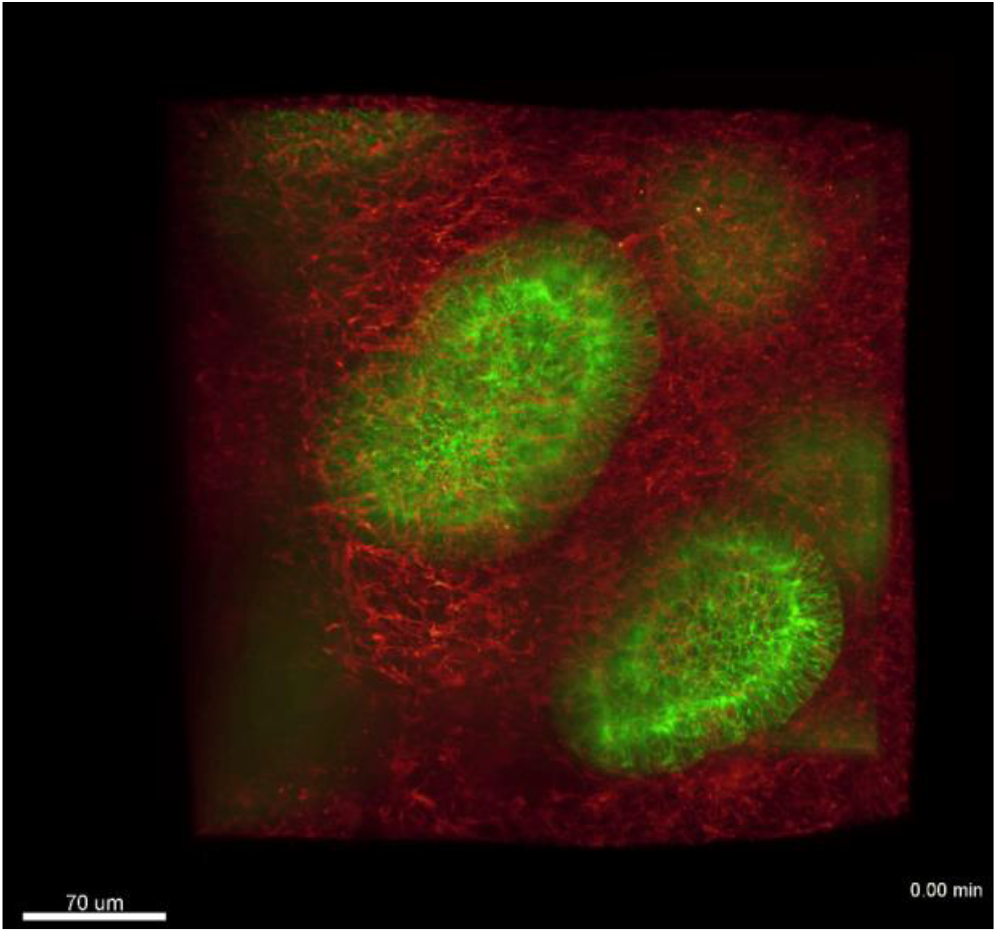
High-resolution light-sheet microscopy timelapse imaging of epithelial lung development (3 hours). Timelapse movie showing the development of an E12.5 mouse lung rudiment carrying the Shh^GC/+^; ROSA^mT/mG^ construct Embryonic lung was mounted in a hollow cylinder made from low-melting-point agarose and filled with matrigel to replicate the native microenvironment and promote near-physiological growth. The sample was imaged using a Zeiss Z.1 Lightsheet, with frames acquired every 20 minutes. This subset includes all 11 time points (3 hours) used in Figure 6. Top bud was used for quantifications. (AVI 59.2 MB).

**Video 8.**
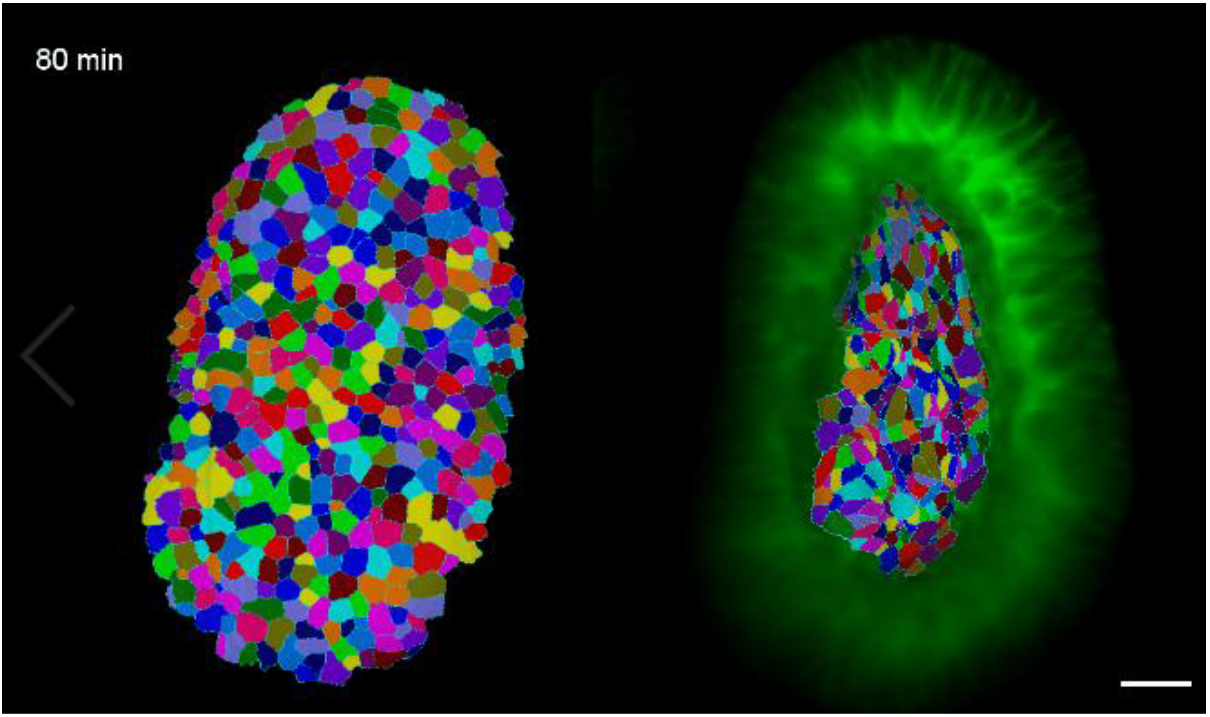
Apical and basal surface cell segmentations. Basal and apical cell segmentations overlays for a CUBIC-cleared E12.5 ShhShh^GC/+^; ROSA^mT/mG^ mouse lung tip imaged using light-sheet microscopy every 20 minutes. MorphoGraphX was used to accurately extract curved surface meshes from 3D volumetric data. The resulting curved (2.5D) surface images of the apical and basal domains were segmented using the Watershed algorithm. Scale bar 20 μm. (AVI 58.1 MB).

**Video 9.**
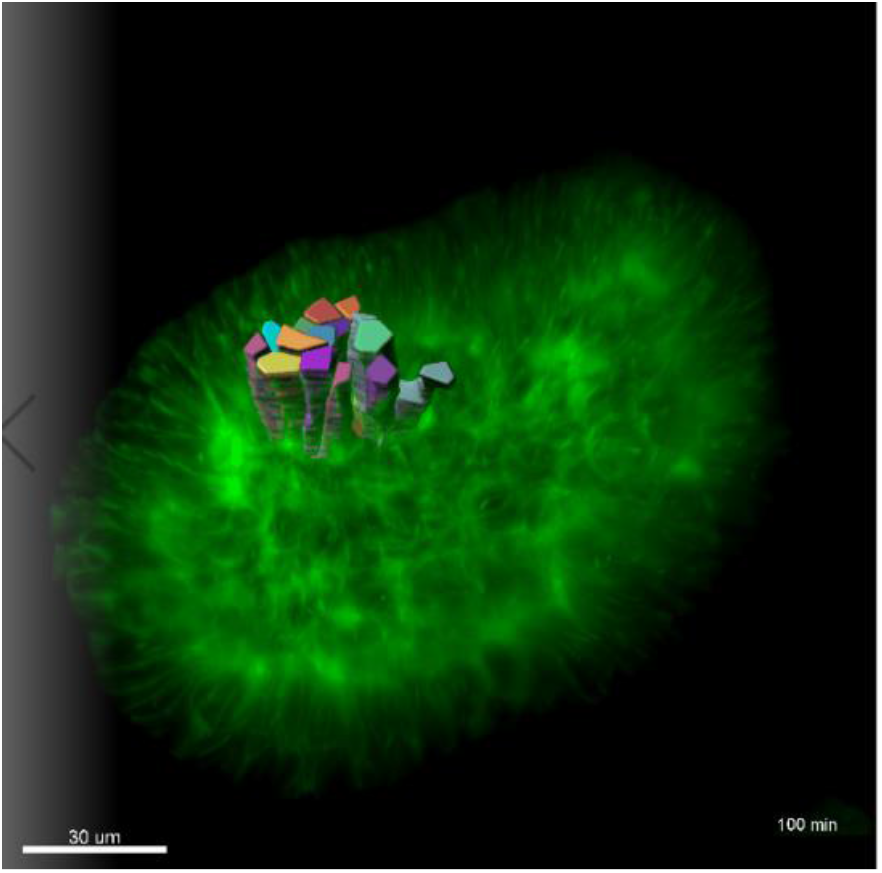
3D timelapse segmentation of growing epithelia. Timelapse segmentation of growing epithelial cells from a CUBIC-cleared E12.5 mouse lung tube over 180 minutes, iso-surfaces colored by cell identifier. This specimen expressed the Shh^GC/+^; ROSA^mT/mG^ reporter and was imaged using light-sheet microscopy. (AVI 9.45 MB).

